# SMC Complexes Are Guarded by the SUMO Protease Ulp2 Against SUMO-Chain-Mediated Turnover

**DOI:** 10.1101/2020.11.13.381483

**Authors:** Ivan Psakhye, Dana Branzei

## Abstract

Structural maintenance of chromosomes (SMC) complexes, cohesin, condensin and Smc5/6, are essential for viability and participate in multiple processes, including sister chromatid cohesion, chromosome condensation, and DNA repair. Here we show that SUMO chains target all three SMC complexes and are antagonized by the SUMO protease Ulp2 to prevent their turnover. We uncover that the essential role of the cohesin-associated subunit Pds5 is to counteract SUMO chains jointly with Ulp2. Importantly, fusion of Ulp2 to kleisin Scc1 supports viability of *PDS5* null cells and protects cohesin from proteasomal degradation mediated by the SUMO-targeted ubiquitin ligase Slx5/Slx8. The lethality of *PDS5* deleted cells can also be bypassed by simultaneous loss of the PCNA unloader, Elg1, and the cohesin releaser, Wpl1, but only when Ulp2 is functional. Condensin and Smc5/6 complex are similarly guarded by Ulp2 against unscheduled SUMO-chain assembly, which we propose to time the availability of SMC complexes on chromatin.

## INTRODUCTION

Posttranslational modification of proteins with the small ubiquitin-like modifier (SUMO) is essential for eukaryotic cells as it regulates substrate fate by affecting protein interactions, activity, localization and abundance (1). SUMOylation frequently targets entire protein groups, actively engaged in common functions (2, 3), and fosters protein complex formation through SUMO binding to SUMO-interacting motifs (SIMs). Similar to ubiquitin, SUMO is conjugated to exposed lysines on the substrates leading to monoSUMOylation or, if several lysines are modified, to multiSUMOylation. Moreover, mono/multiSUMOylation may be extended to SUMO chains when lysines of SUMO conjugated to the substrate are being further modified with SUMO, leading to substrate polySUMOylation. The modifications are reversible and counteracted by SUMO proteases, which have different localization, substrate and SUMO-linkage specificities (4). In budding yeast, the SUMO protease Ulp2 has preference for SUMO chains and prevents substrate polySUMOylation, which can be further recognized by ubiquitin E3 ligases containing multiple SIMs. These so-called SUMO-targeted ubiquitin ligases (STUbLs) can mediate proteolytic or non-proteolytic ubiquitylation of the SUMOylated substrate (5). Thus, SUMO chains disassembled by the Ulp2 protease may function as a countdown timer if they are assembled on the substrates of STUbLs.

We recently reported that SUMO-chain/Ulp2-protease-regulated proteasomal degradation is a mechanism that times the availability of DDK, a key replication initiation regulator (6). To extend our findings beyond DNA replication, we performed an unbiased SILAC-based proteomic screen to uncover degradation-prone SUMO conjugates that decrease in abundance in the absence of Ulp2 specifically in a SUMO-chain-dependent manner. Strikingly, we found subunits of all three SMC complexes, cohesin, condensin and Smc5/6 (7, 8) as potential hits, suggesting that the abundance of SMC complexes is regulated via a SUMO-chain-dependent mechanism.

Cohesin was previously shown to become SUMOylated upon loading onto DNA, and loss of cohesin SUMOylation resulted in cohesion defects (9, 10). Moreover, the cohesin associated factor, Pds5, was proposed to protect monoSUMOylated cohesin and facilitate cohesion by preventing via a yet unidentified mechanism Siz2 SUMO-ligase-mediated cohesin polySUMOylation that leads to increased Slx5/8 STUbL-mediated proteasome degradation of the cohesin kleisin, Scc1 (11). More recently, depletion of *SENP6,* the human ortholog of Ulp2, was shown to decrease cohesin binding to chromatin and cause cohesion defects (12). Taken together, these data suggest that mono/multiSUMOylation of DNA-loaded cohesin is important for cohesion, whereas polySUMOylation induced in *pds5* mutants targets cohesin for STUbL-mediated proteasomal turnover compromising cohesion.

Here we aimed to address if and how Ulp2 protects cohesin and other SMC complexes from SUMO-chain-targeted turnover. We find that fusion of Ulp2 to the cohesin’s kleisin Scc1 protects cohesin from proteasomal degradation and supports viability in the complete absence of Pds5 that is otherwise essential. Moreover, we identify that simultaneous loss of the PCNA unloader, Elg1, and the cohesin releaser, Wpl1, allows viability of *PDS5* null cells in a manner strictly depending on Ulp2 function. These results indicate that the essential function of Pds5 is to counteract SUMO chains jointly with Ulp2 rather than support cohesin activity in other direct ways. Condensin and Smc5/6 are also safeguarded by Ulp2 against unscheduled SUMO chain assembly, overall suggesting a SUMO-chain/Ulp2-protease-governed mechanism that instructs SMC complexes availability on chromatin.

## MATERIAL AND METHODS

### Yeast Strains

Chromosomally tagged *Saccharomyces cerevisiae* strains and mutants were constructed by a PCR-based strategy, by genetic crosses and standard techniques (13). Standard cloning and site-directed mutagenesis techniques were used. Strains and all genetic manipulations were verified by polymerase chain reaction (PCR), sequencing and phenotype. All yeast strains used in this work except those specifically indicated and used for the yeast two-hybrid (Y2H) studies are isogenic to W303 background and are listed in the Table 1.

**Table 1.**
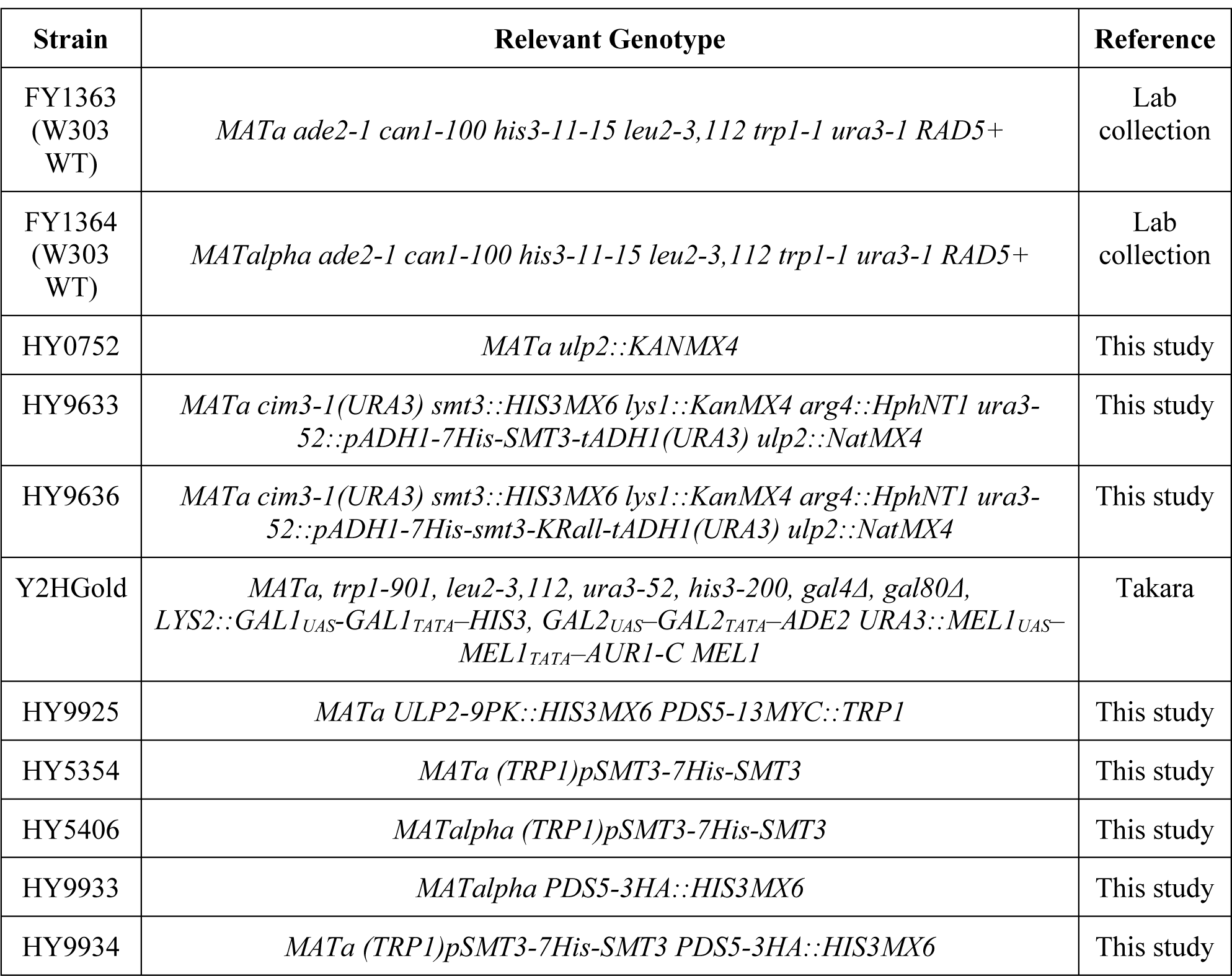
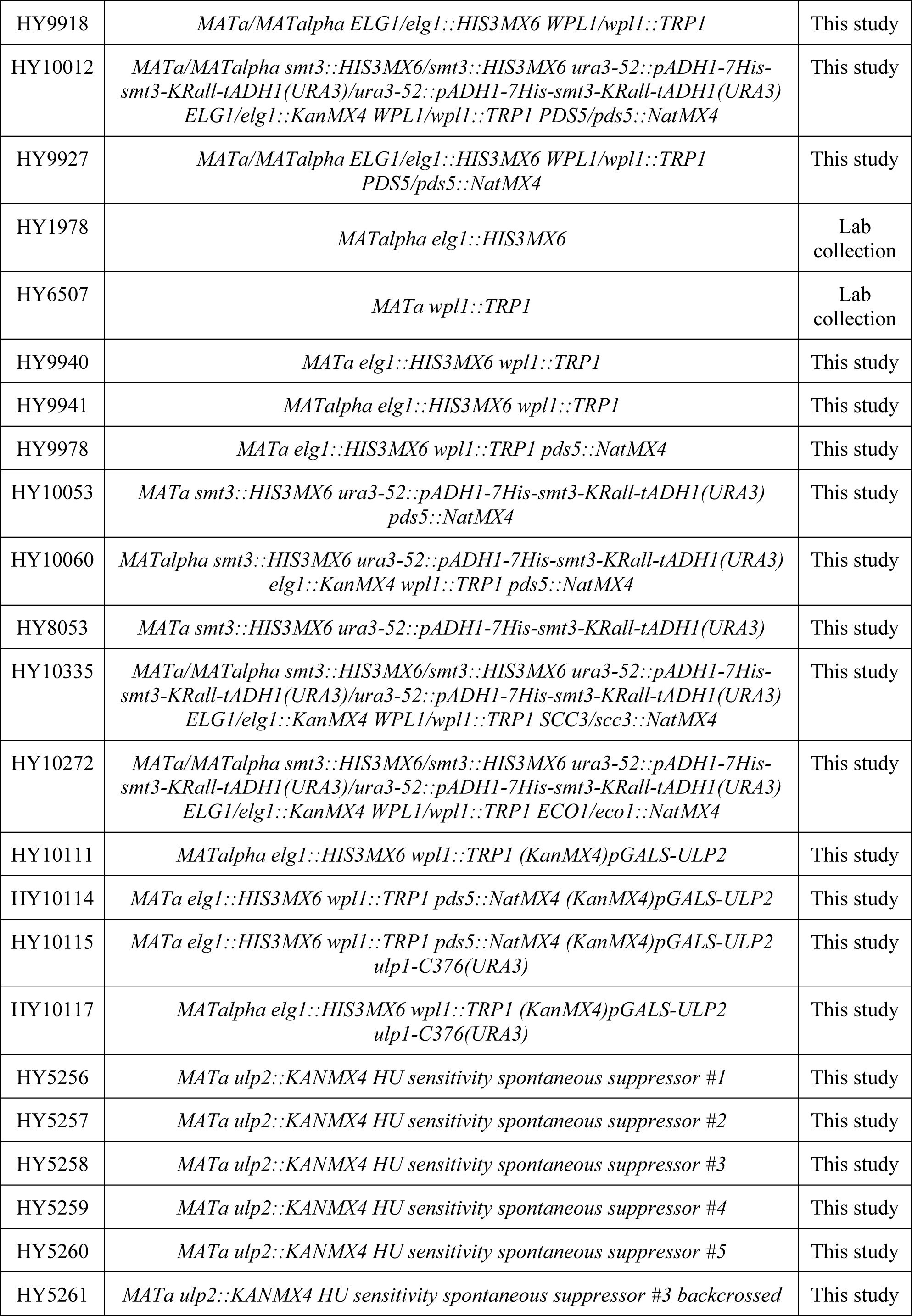
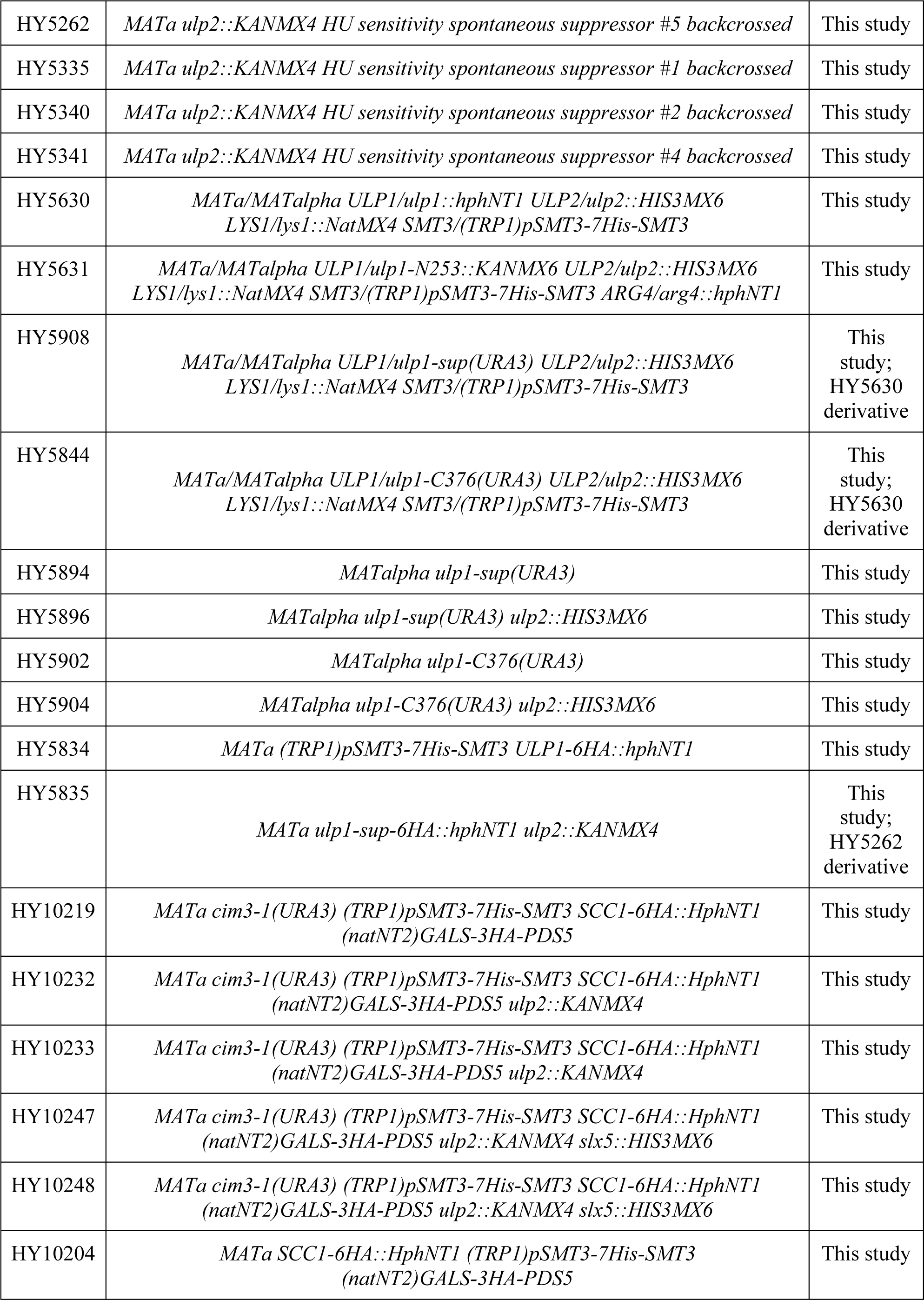
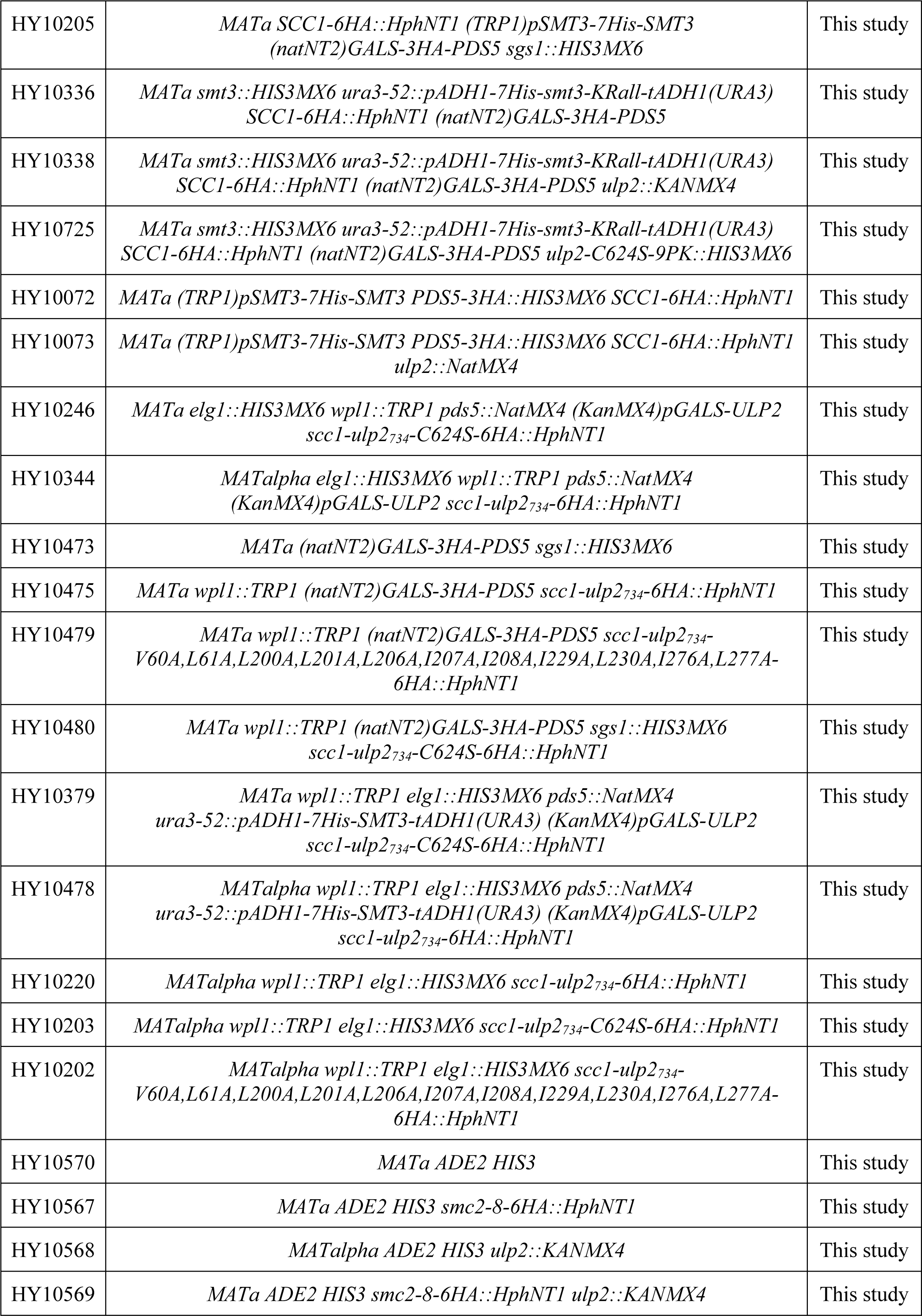
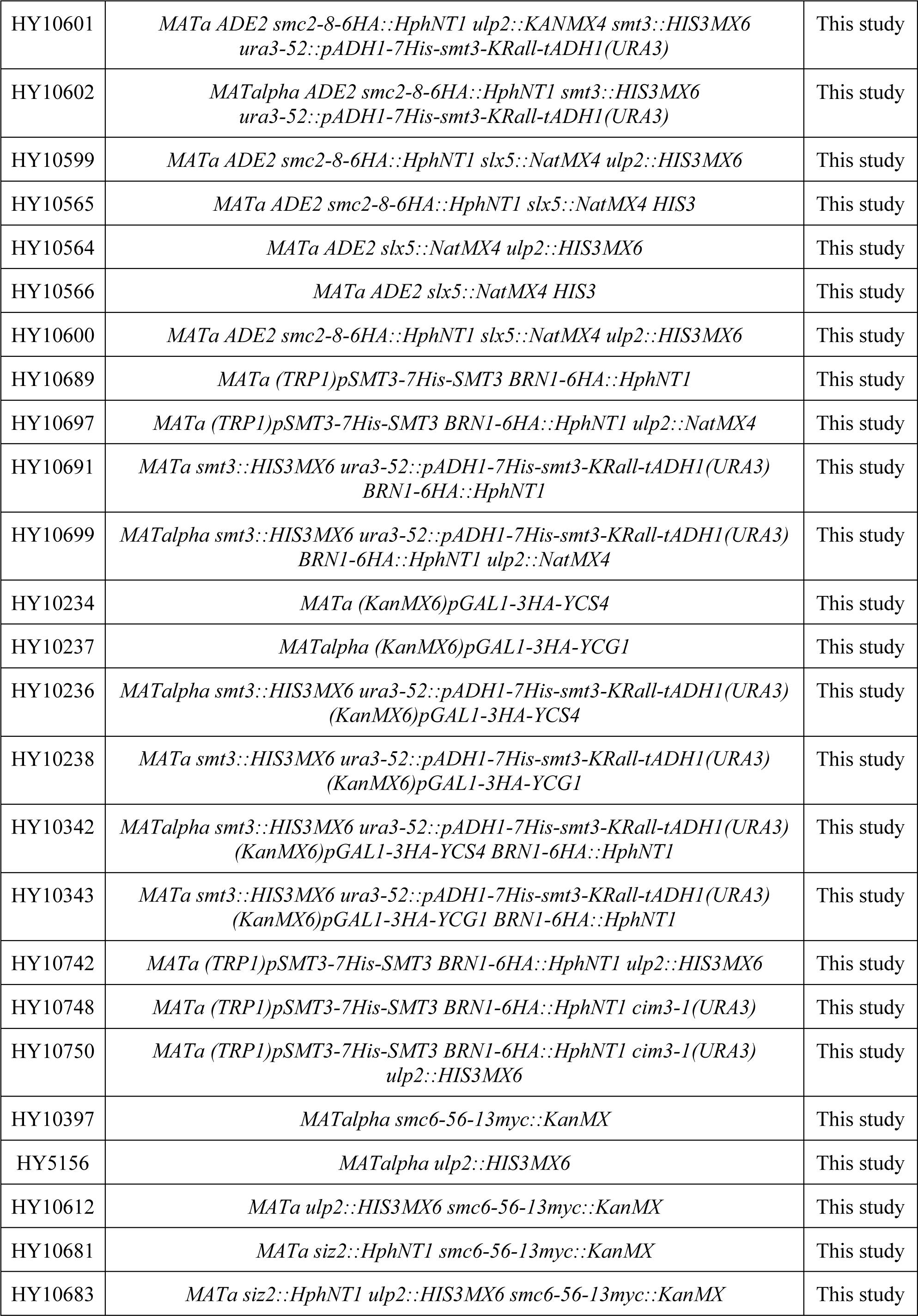
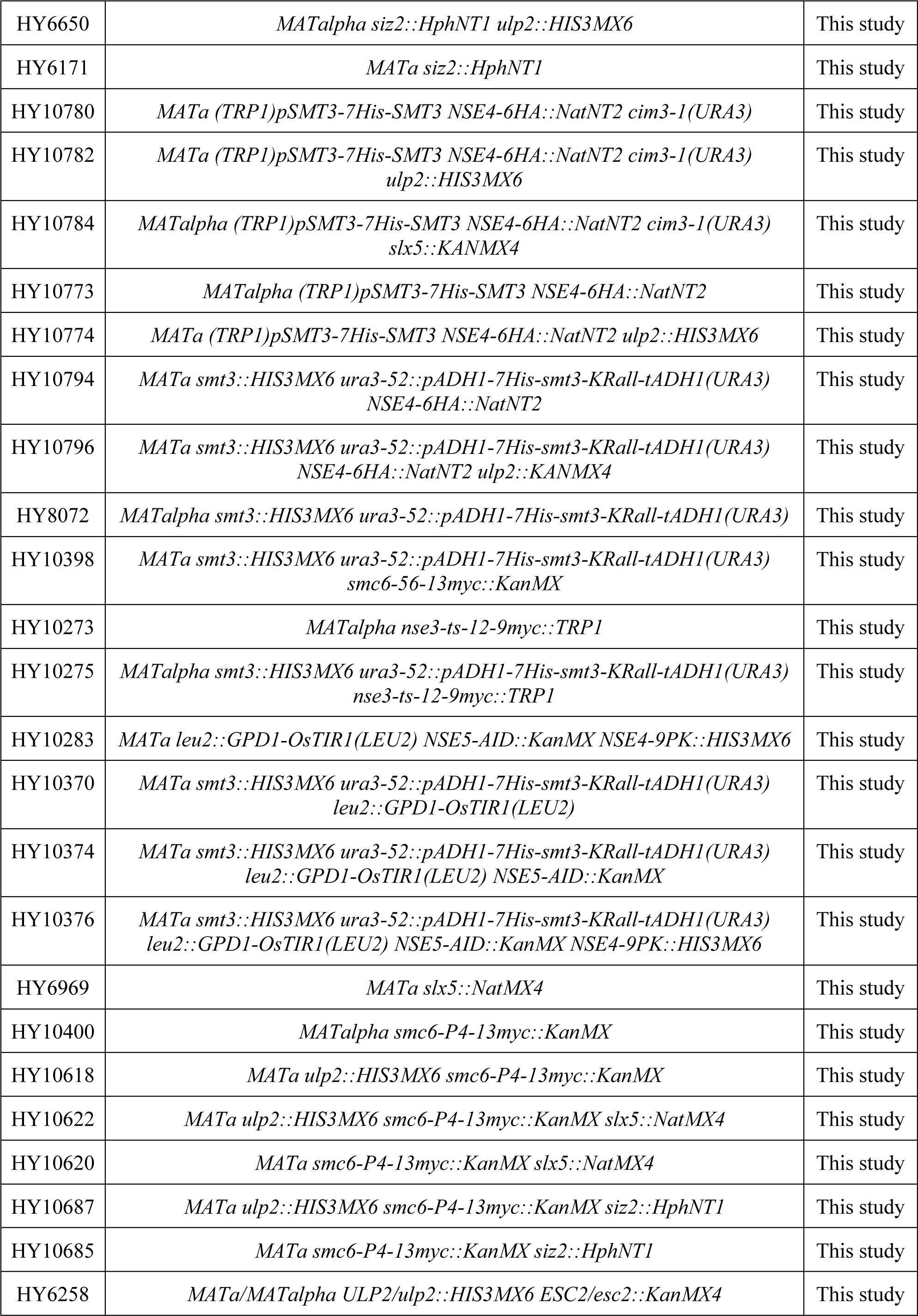
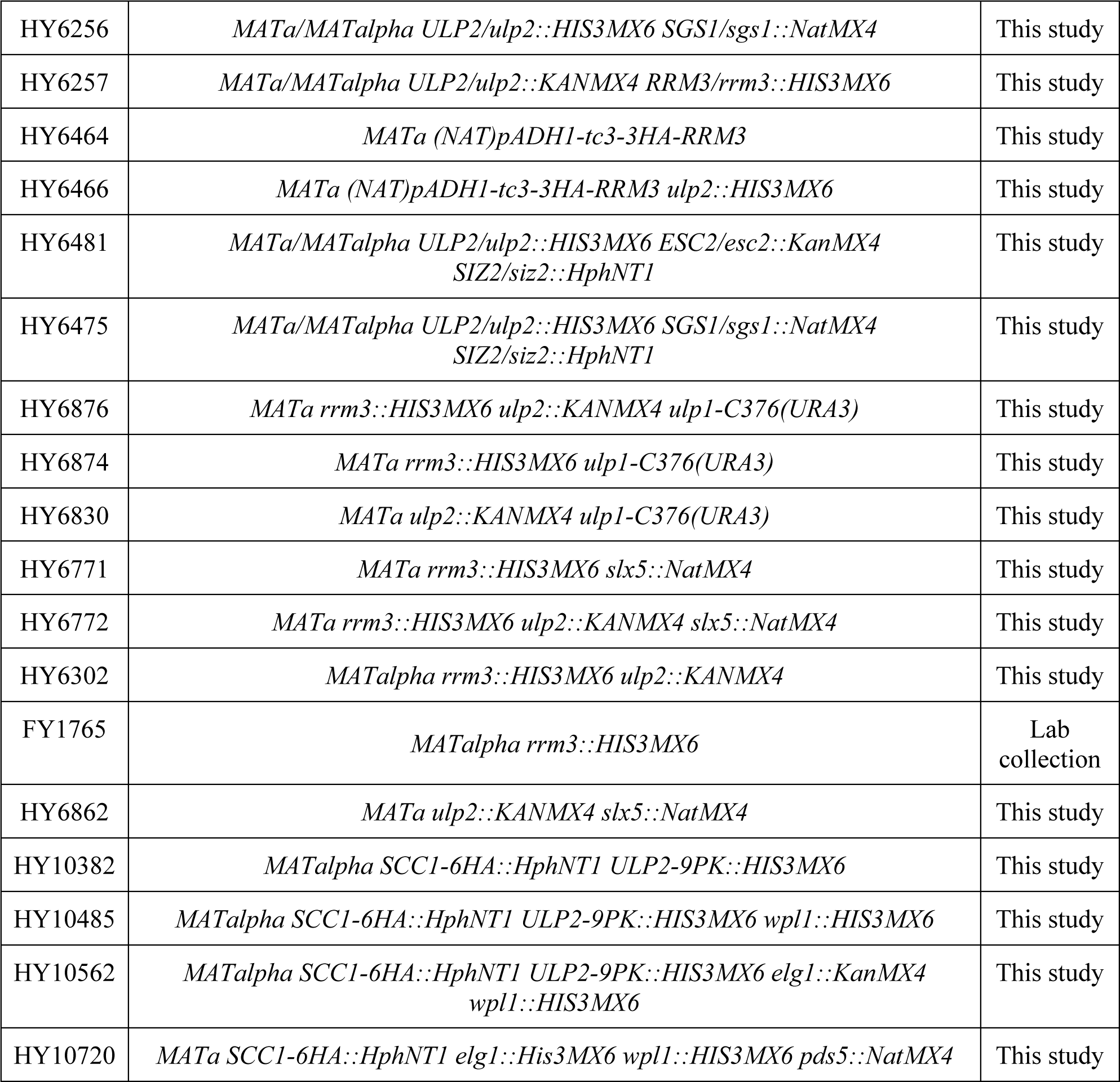
Saccharomyces cerevisiae strains used in this study.

All strains are isogenic to W303 background, except for HY10570, HY10567, HY10568, HY10569, HY10600, HY10601, HY10602, HY10599, HY10565, HY10564, HY10566, which are derived from VG2029-7B (14), and Y2HGold (Takara).

### Yeast Techniques

Yeast cultures were inoculated from overnight cultures, grown using standard growth conditions and media (15). All cultures were grown in YPD media containing glucose (2%) as carbon source at 28°C unless otherwise indicated. For the transcriptional shut-off of genes expressed under the control of *GAL* promoter, cells were grown in YP Gal media containing galactose (2%), washed once with 1X PBS and shifted to YPD media or plated on YPD plates. For drug sensitivity assays, cells from overnight cultures were counted and diluted before being spotted on YPD plates containing the indicated concentrations of drugs and incubated at 28°C for 2-3 days. For Y2H analysis catalytically-dead *ulp2-S624S* mutant and different truncations of *PDS5* were cloned into pGAD-C1 or pGBD-C1 vectors and cotransformed into Y2HGold yeast strain. Standard cloning and site-directed mutagenesis techniques were used. Maps and primer DNA sequences are available upon request.

### TCA Protein Precipitation

To preserve the post-translational modifications, yeast cells were lysed under denaturing conditions. For preparation of denatured protein extracts, yeast cultures grown to an OD_600_=0.7-1 were pelleted by centrifugation (4000 rpm, 4 min, 4°C) and immediately frozen in liquid nitrogen. After thawing on ice, the pellets were lysed by addition of denaturing lysis buffer (1.85 M NaOH, 7.5% β-mercaptoethanol) for 15min on ice. For the cell pellet of an OD_600_=1 typically 150 μl of lysis buffer was used. To precipitate the proteins, the lysate was subsequently mixed with an equal volume (150 μl in case of OD_600_=1) of 55% (w/v) trichloroacetic acid (TCA) and further incubated on ice for 15 min. The precipitated material was recovered by two sequential centrifugation steps (13000 rpm, 4°C, 15 min). Pelleted denatured proteins were then either directly resuspended in HU sample buffer (8 M urea, 5% SDS, 1 mM EDTA, 1,5% DTT, 1% bromophenol blue; 50 μl per OD_600_=1), boiled for 10 min and stored at -20°C, or used for downstream processing, e.g., Ni-NTA pull-downs of His-tagged SUMO conjugates.

### Ni-NTA Pull-down of ^His^SUMO Conjugates

For isolation of *in vivo* SUMOylated substrates from yeast cells expressing N-terminally His-tagged Smt3 (^His^SUMO), denatured protein extracts were prepared and Ni-NTA chromatography was carried out as described previously (2, 16). In general, 200 OD_600_=1 of logarithmically growing cells were harvested by centrifugation (4000 rpm, 4 min, 4°C), washed with pre-chilled water, transferred to 50 ml falcon tube and lysed with 6 ml of 1.85 M NaOH / 7.5% β-mercaptoethanol for 15 min on ice. The proteins were precipitated by adding 6 ml of 55% TCA and another 15 min incubation on ice (TCA-precipitation, described above). Next, the precipitate was pelleted by centrifugation (3500 rpm, 15 min, 4°C), washed twice with water and finally resuspended in buffer A (6 M guanidine hydrochloride, 100 mM NaH2PO4, 10 mM Tris-HCl, pH 8.0, 20 mM imidazole) containing 0.05% Tween-20. After incubation for 1 hour on a roller at room temperature with subsequent removal of insoluble aggregates by centrifugation (23000 g, 20 min, 4°C), the protein solution was incubated overnight at 4°C with 50 μl of Ni-NTA agarose beads in the presence of 20 mM imidazole. After incubation, the beads were washed three times with buffer A containing 0.05% Tween-20 and five times with buffer B (8 M urea, 100 mM NaH2PO4, 10 mM Tris-HCl, pH 6.3) with 0.05% Tween-20. ^His^SUMO conjugates bound to the beads were finally eluted by incubation with 50 μl of HU sample buffer for 10 min at 65°C. Proteins were resolved on precast Bolt 4%–12% Bis-Tris Plus gradient gels, and analyzed by standard Western blotting techniques using antibodies listed below.

### Antibodies

Mouse monoclonal anti-Viral V5-TAG antibody (1:5000; clone SV5-Pk1) was purchased from Bio-Rad/AbD Serotec. Mouse monoclonal anti-Pgk1 antibody (1:2000; clone 22C5D8) was obtained from Thermo Fisher Scientific. Mouse monoclonal anti-HA antibody (1:2000; clone F-7) and rabbit polyclonal anti-Smt3 (1:2000; clone y-84) antibody were from Santa Cruz Biotechnology, as well as normal mouse IgG. Mouse monoclonal anti c-MYC antibody (1:2000; clone 9E10) was produced in house. Mouse monoclonal anti-acetyl-Smc3 antibody (17) (1:2000) was a kind gift from Katsuhiko Shirahige. Anti-rabbit IgG and anti-mouse IgG, HRP-linked antibodies (1:5000) were purchased from Cell Signaling Technology.

### Immunoprecipitation

For the immunoprecipitation (IP) and binding studies involving co-IP, native yeast extracts were prepared by cell disruption using grinding in liquid nitrogen. To avoid protein degradation and loss of PTMs, lysis buffer (150 mM NaCl, 10% glycerol, 1% NP-40, 50 mM Tris HCl, pH 8.0) was supplemented with inhibitors: EDTA-free complete cocktail, 20mM N-ethylmaleimide, 1 mM phenylmethanesulfonyl fluoride (PMSF), 25 mM iodoacetamide, and phosphatase inhibitor cocktails 2 and 3 (Sigma-Aldrich). For IPs, anti-PK and anti-MYC antibodies, together with recombinant protein G Sepharose 4B beads were used. IPs were performed overnight with head-over-tail rotation at 4°C and were followed by stringent washing steps to remove non-specific background binding to the beads.

### ChIP-qPCR

Chromatin immunoprecipitation (ChIP) was carried out as previously described (6). Briefly, cells were collected at the indicated experimental conditions and crosslinked with 1% formaldehyde for 30 min. Cells were washed twice with ice-cold 1X TBS, suspended in lysis buffer supplemented with 1 mM PMSF, 20 mM NEM, and 1X EDTA-free complete cocktail, and lysed using FastPrep-24 (MP Biomedicals). Chromatin was sheared to a size of 300-500 bp by sonication. IP reactions with anti-HA antibodies and Dynabeads protein G were allowed to proceed overnight at 4°C. After washing and eluting the ChIP fractions from beads, crosslinks were reversed at 65°C overnight for both Input and IP. After proteinase K treatment, DNA was extracted twice by phenol/chlorophorm/isoamyl alcohol (25:24:1, v/v). Following precipitation with ethanol and Ribonuclease A (RNase A) treatment, DNA was purified using QIAquick PCR purification kit. Real-time PCR was performed using QuantiFast SYBR Green PCR kit according to the manufacturer’s instructions and each reaction was performed in triplicates using a Roche LightCycler 96 system. The results were analyzed with absolute quantification/2^nd^ derivative maximum and the 2(-ΔC(t)) method. Each ChIP experiment was repeated at least three times. Statistical analysis was performed using Student’s unpaired *t*-test. The error bars represent standard error of mean (SEM).

### Mass Spectrometry

For the detection of degradation-prone SUMO conjugates decreased in abundance in *ulp2*Δ *cim3-1* mutant cells specifically in a SUMO-chain-dependent manner (Figure 1), SILAC-based mass spectrometry protocol (18) was used. Yeast *ulp2*Δ *cim3-1* mutant cells deficient in biosynthesis of lysine and arginine (*lys1*Δ and *arg4*Δ) expressing either wild-type His-tagged SUMO (^His^SUMO) or its lysine-less variant (*KRall*) that cannot form lysine-linked polySUMO chains were grown for at least ten divisions in synthetic complete media supplemented either with unlabeled (Lys0 and Arg0; light) or heavy isotope-labeled amino acids (Lys8 and Arg10; heavy) from Cambridge Isotope Laboratories. Exponentially dividing *^His^SUMO ulp2*Δ *cim3-1* cells grown in heavy media were harvested, combined with equal amount of *KRall ulp2*Δ *cim3-1* cells grown in light media, and SUMO conjugates were isolated by using denaturing Ni-NTA pull-down. Proteins isolated following denaturing Ni-NTA pull-downs of ^His^SUMO conjugates were separated on 4-12% Bis-Tris gel. The whole lane was excised in slices and proteins were digested with trypsin. Extracted peptides were analyzed by LC-MS/MS using the Q Exactive HF mass spectrometer and identified using MaxQuant (19) software.

**Figure 1.**
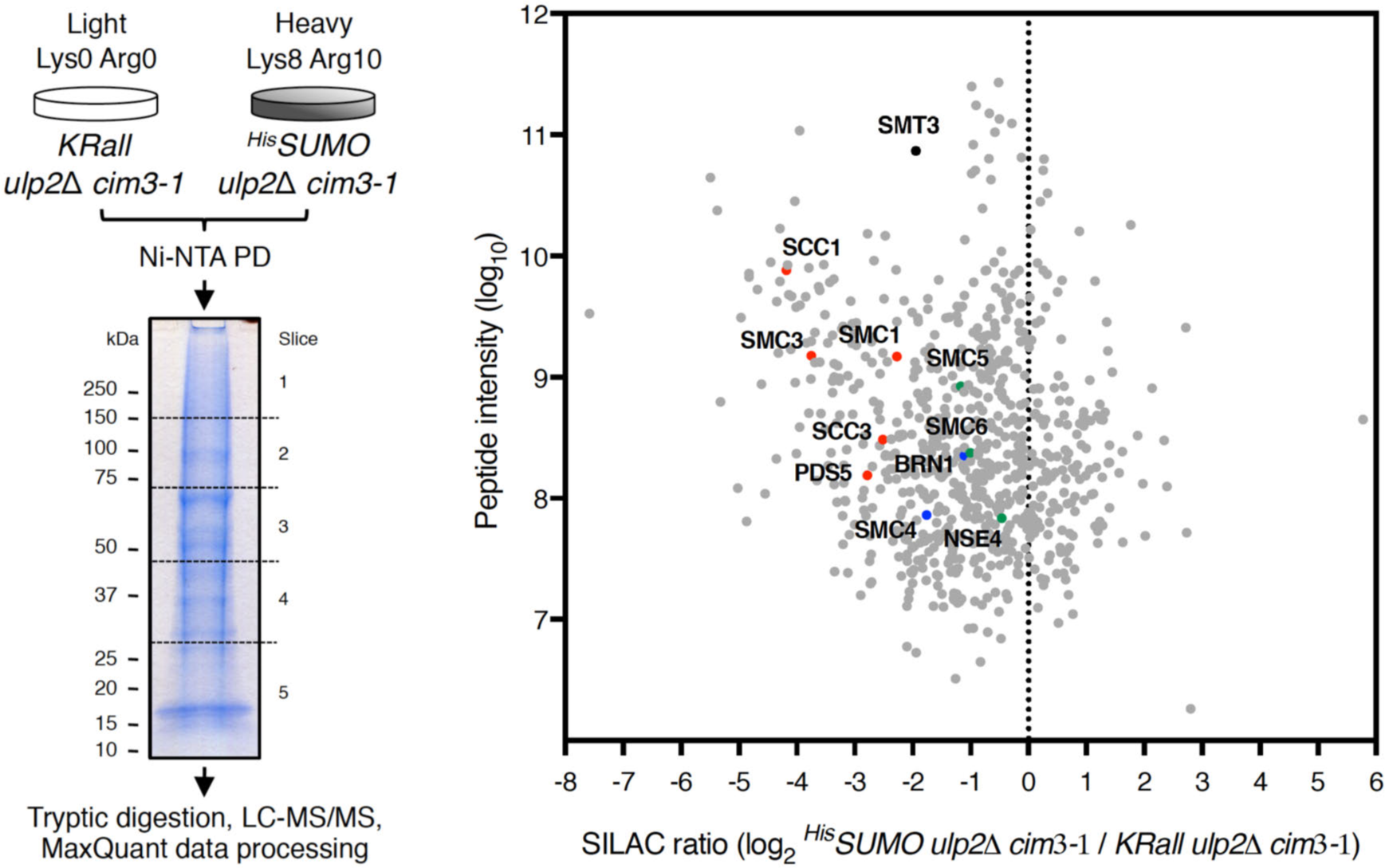
SUMO chains promote the turnover of SUMOylated SMC complex subunits in the absence of Ulp2 SUMO protease. Outline of SILAC experiment performed to detect degradation-prone SUMOylated substrates that decrease in abundance in a SUMO-chain-dependent manner in *ulp2*Δ *cim3-1* cells (left). SILAC ratios for 726 quantified proteins plotted against the sum of the relevant peptide intensities (right). SUMOylated subunits of cohesin (red), condensin (blue) and the Smc5/6 complex (green) accumulate if instead of ^His^SUMO, which is able to form SUMO chains, a lysine-less SUMO variant (*KRall*) that cannot form lysine-linked SUMO chains is expressed as the only source of SUMO.

## RESULTS

### SUMO chains target SMC complexes and promote their turnover

We have used quantitative proteomics to identify SUMO substrates whose turnover is promoted by SUMO chains in the absence of the SUMO protease Ulp2 that possesses SUMO-chain-editing activity in yeast cells (20). To these ends, we used a SILAC-based mass spectrometry approach (18) and compared by denaturing Ni-NTA pull-down (Ni PD (2, 16)) the levels of SUMO conjugates in *ulp2*Δ *cim3-1* mutants expressing either endogenous yeast SUMO (*SMT3*) N-terminally tagged with a 7xHis-tag (^His^SUMO) or a lysine-less SUMO variant (*KRall*) that cannot form lysine-linked polySUMO chains (Figure 1, left). We used the temperature sensitive proteasome defective *cim3-1* mutant cells at the permissive temperature of 30°C to allow cell cycle progression and accumulation of degradation-prone substrates, thus facilitating their identification by mass spectrometry. The SILAC screen quantified 726 potential SUMO conjugates (Figure 1, right); the abundance of most of them did not change significantly, while SUMO conjugates (Smt3) pulled-down from the *KRall* mutant were more abundant in general. Notably, among the SUMO substrates strongly enriched in the sample derived from chainless SUMO *ulp2*Δ *cim3-1* cells, were subunits of all three SMC complexes. SUMOylated cohesin subunits Smc1, Smc3, Scc1, Scc3, and Pds5 were enriched the most when SUMO-chain growth was prevented by the *KRall* mutation, whereas SUMOylated condensin subunits Smc4, Brn1, and the Smc5/6 complex subunits Smc5, Smc6, Nse4 accumulated to a lesser extent.

Cohesin shows strong ties to the SUMO system (9-11,21-23). Its regulatory subunit Pds5 is a known SUMO target ((14) and Supplementary Figure S1A) and is one of the first identified substrates of Ulp2 (14). These findings, together with the results of our SILAC screen suggesting that most of cohesin subunits are subjected to SUMO-chain-mediated turnover, prompted us to focus on studying the regulation of the cohesin complex by SUMO chains and Ulp2.

We hypothesized that one of the roles of the essential regulatory cohesin subunit Pds5 is to recruit Ulp2 to protect cohesin against unscheduled SUMO-chain-mediated turnover. Indeed, we could confirm the interaction of Ulp2 with Pds5 using both co-immunoprecipitation (co-IP) and yeast two-hybrid (Y2H) studies (Supplementary Figure S1B-D). For the co-IP, we C-terminally tagged endogenous Pds5 and Ulp2 with 13Myc and 9PK tags, respectively. IPs with the anti-PK (Supplementary Figure S1B) and anti-Myc (Supplementary Figure S1C) antibodies revealed that Ulp2 interacts with Pds5 and there is preference towards upshifted, potentially SUMO-modified forms of proteins. These slower-migrating species of Pds5 and Ulp2 were co-immunoprecipitated with specific antibodies, but not mouse IgG (Supplementary Figure S1B-C). For the Y2H studies, we used Gal4 DNA-binding domain (BD) fusions of various Pds5 truncations and the Gal4 activation domain (AD) fusion of catalytically dead Ulp2 (Ulp2-C624S; Ulp2CD), which we expected to interact stronger with potential substrates based on previous work on Ulp1CD that behaved like a SUMO substrate trap (24) (Supplementary Figure S1D-F). We observed weak Y2H interaction of Ulp2 with both N-terminal (aa 1-701) and C-terminal (aa 702-1277) fragments of Pds5 (Supplementary Figure S1D). However, analysis of further N-terminal Pds5 truncations revealed auto-activation of the *HIS3* reporter gene by Pds5 N-terminus (aa 1-250, Supplementary Figure S1E), suggesting that binding of Ulp2 to Pds5 may be mediated by its C-terminus. Indeed, using C-terminal truncations of Pds5 we found that the C-terminal fragment of Pds5 (aa 1078-1277) is required for the interaction with Ulp2 in the Y2H system (Supplementary Figure S1F). Altogether, these results potentially support the notion that Pds5 recruits Ulp2 to prevent SUMO chain-mediated turnover of cohesin.

### The essential role of Pds5 relates to curbing down SUMO chains

Cohesin plays critical roles in numerous cellular pathways (8,25–27), including sister chromatid cohesion and chromosome condensation, for both of which *PDS5* is required (28, 29). Interestingly, sister chromatid cohesion defects and lethality of the temperature sensitive *pds5-1* mutant are suppressed by deletion of *ELG1* (30), which acts in the context of the Elg1-Rfc2-5 complex as principal unloader of chromatin-bound proliferating cell nuclear antigen (PCNA) (31). Chromosome condensation defects of *pds5-1* mutant, but not lethality, are in turn suppressed by deletion of *RAD61* (*WPL1*) (30), which encodes the cohesin release factor WAPL that interacts with Pds5 and destabilizes cohesin’s binding to DNA (32).

To study the potential role of Pds5 in counteracting SUMO chains via Ulp2 recruitment to cohesin, we decided to examine the effect of *elg1*Δ and *wpl1*Δ mutations on the viability of *PDS5* null cells when SUMO chain formation is prevented. First, we confirmed that *elg1*Δ *wpl1*Δ double mutant does not affect cell growth compared to single mutants and WT cells (Figure 2A and (33)). Then, we generated *ELG1*/*elg1*Δ *WPL1*/*wpl1*Δ *PDS5*/*pds5*Δ *smt3-KRall*/*smt3-KRall* diploid strain expressing chainless SUMO variant *smt3-KRall* as the only source of SUMO. The analysis of resulting haploids after sporulation and tetrad dissection of this strain should reveal if *elg1*Δ *wpl1*Δ *smt3-KRall* triple mutant bypasses the essential role(s) of *PDS5*. Strikingly, we found that not only *pds5*Δ *elg1*Δ *wpl1*Δ *smt3-KRall* quadruple mutant was viable, but that expression of *smt3-KRall* alone bypassed the requirement of *PDS5* for viability (Figure 2B). We next analyzed if *elg1*Δ *wpl1*Δ *pds5*Δ triple mutant is viable following tetrad dissection of *ELG1*/*elg1*Δ *WPL1*/*wpl1*Δ *PDS5*/*pds5*Δ diploid strain. Interestingly, also the *elg1*Δ *wpl1*Δ double mutant suppressed the lethality of *pds5*Δ cells (Figure 2C). Thus, we uncovered that the essential role of *PDS5* in budding yeast is linked to counteracting SUMO chains and identified a genetic background, *elg1*Δ *wpl1*Δ, in which this essential function is bypassed.

**Figure 2.**
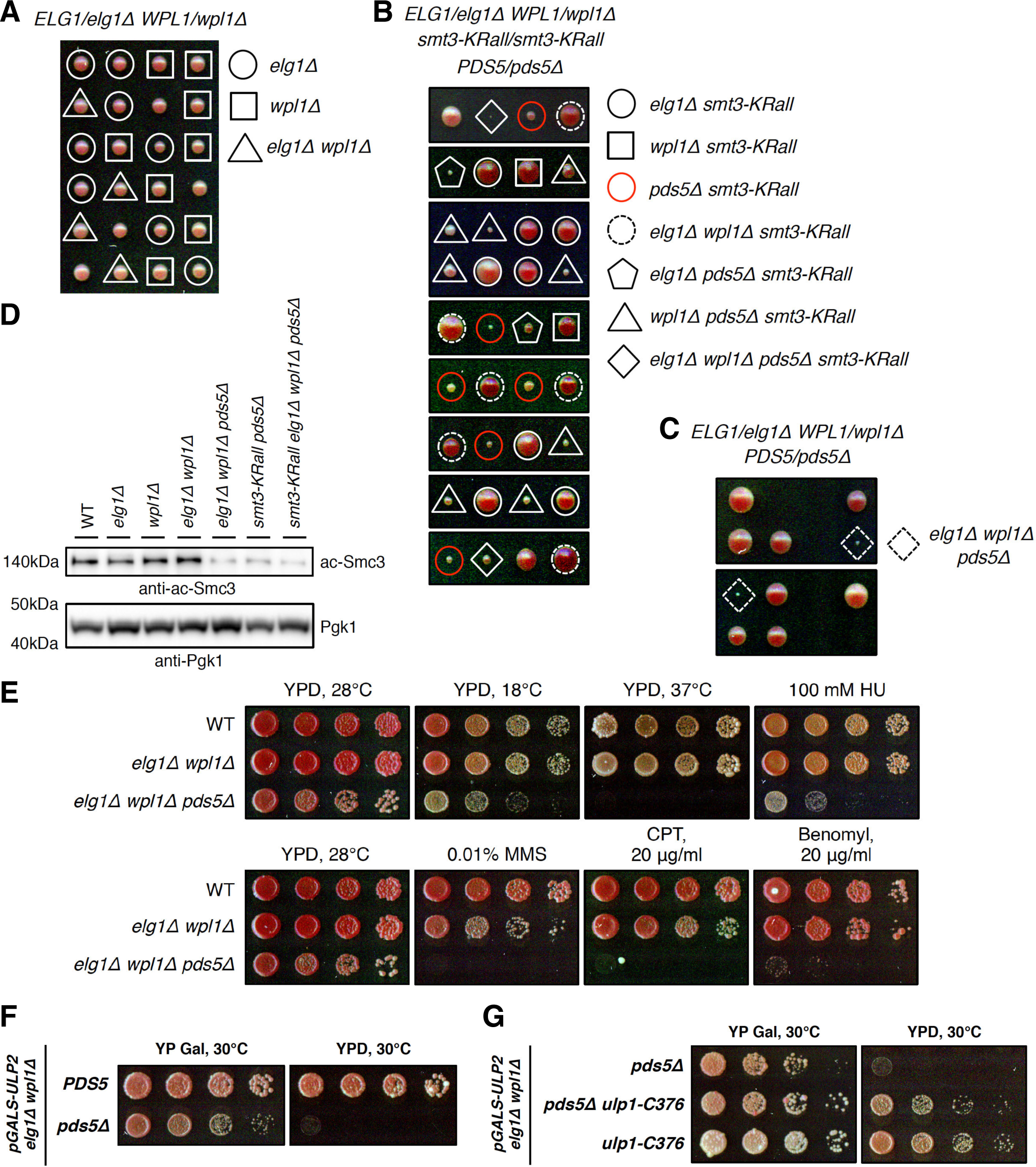
The essential role of cohesin-associated subunit Pds5 is to counteract SUMO chains and is bypassed in the *elg1*Δ *wpl1*Δ background. (A) Tetrad dissection analysis of *ELG1*/*elg1*Δ *WPL1*/*wpl1*Δ double mutant revealed no negative genetic interaction compared to WT. (B) Lethality of *pds5*Δ cells is suppressed by SUMO mutant *smt3-KRall* that cannot form lysine-linked SUMO chains. (C) Lethality of *pds5*Δ cells is bypassed in *elg1*Δ *wpl1*Δ background. (D) Smc3 lysine K112, K113 acetylation is decreased in *pds5*Δ cells, lethality of which is suppressed either by *smt3-KRall, elg1*Δ *wpl1*Δ, or their combination. (E) *pds5*Δ *elg1*Δ *wpl1*Δ cells are slow-growth and exhibit sensitivity to temperature, HU, MMS, CPT and benomyl. Spotting of 1:7 serial dilutions on YPD plates at indicated temperatures and drug concentrations. (F) Viability of *pds5*Δ *elg1*Δ *wpl1*Δ cells depends on *ULP2*. Upon transcriptional shut-off of *ULP2* expressed from inducible *GAL* promoter *pds5*Δ *elg1*Δ *wpl1*Δ cells die. (G) Lethality of *pds5*Δ *elg1*Δ *wpl1*Δ *ulp2* cells is bypassed by the spontaneous suppressor of *ulp2*Δ phenotypes denoted *ulp1-C376* that is no longer tethered to the nuclear pores and is able to deSUMOylate Ulp2 substrates in the nucleoplasm.

Next, we checked if expression of the chainless SUMO variant *smt3-KRall, elg1*Δ *wpl1*Δ double mutant, or their combination, is able to suppress the loss of another cohesin HAWK protein Scc3, which is required for cohesin binding to DNA (34), but not for its loading to chromatin mediated by Scc2/Scc4 complex (35). None of the mutations provided viability to *scc3*Δ haploids upon tetrad dissection of *ELG1*/*elg1*Δ *WPL1*/*wpl1*Δ *SCC3*/scc3Δ *smt3-KRall*/*smt3-KRall* diploid strain (Supplementary Figure S2A). The lethality of cells upon deletion of *ECO1*, the acetyltransferase required for Smc3 lysine K112, K113 acetylation and sister chromatid cohesion establishment (36, 37), is also not suppressed by expressing *smt3-KRall* and is only bypassed by *wpl1*Δ (Supplementary Figure S2B). Thus, while *PDS5* overexpression partially suppresses the temperature sensitivity of certain *eco1* thermosensitive mutants (38), *smt3-KRall* that bypasses the essential role of *PDS5* does not restore viability of cells lacking Eco1 (Supplementary Figure S2B), suggesting that other functions of *PDS5* are required for this suppression.

To determine the consequences of *PDS5* loss for the chromatin-bound cohesin levels, we assessed Eco1-mediated Smc3 lysine K112, K113 acetylation in the identified *PDS5* null bypass conditions. Eco1 acetyltransferase targets cohesin loaded onto DNA (39), thus making Smc3 acetylation a good indicator of the functionally-engaged cohesin amounts operating in the cell. To this end, we utilized a monoclonal antibody specific for the Eco1-mediated Smc3 lysine K112, K113 acetylation (17) and found that it was largely reduced, but not abolished, in all *pds5*Δ mutants (Figure 2D and Supplementary Figure S2C). Thus, cells lacking Pds5 have reduced levels of chromatin-bound cohesin available to fulfil its functions, which is in agreement with findings in *pds5-ts* cells (11, 29). We further found that while limited amounts of chromatin-bound acetylated cohesin in *pds5*Δ *elg1*Δ *wpl1*Δ mutants are sufficient to support viability, the mutant cells are sensitive to low and high temperatures, replication stress induced by hydroxyurea (HU), exposures to the DNA-alkylating agent methyl methanesulfonate (MMS), topoisomerase poison camptothecin (CPT) and microtubule-depolymerizing drug benomyl (Figure 2E).

Loss of *WPL1* was previously reported to cause an increase in pericentromeric cohesin in cells blocked in late G1 by the Cdk1 inhibitor Sic1, while *wpl1*Δ had little or no effect on the extent of Scc1 association with chromosome arms (40). To provide insights on how *elg1*Δ *wpl1*Δ mutant suppresses the lethality of *pds5*Δ cells, we next analyzed the cohesin levels on chromatin at the pericentromeric region of *CEN10*, *TER1004*, and the centromere-distal region on chromosome 3, *ARS305*, by performing chromatin immunoprecipitation (ChIP) of C-terminally 6HA-tagged Scc1 from nocodazole-arrested WT, *wpl1*Δ, *elg1*Δ *wpl1*Δ and *elg1*Δ *wpl1*Δ *pds5*Δ cells (Supplementary Figure S2D-E). Interestingly, we observed statistically significant increase of Scc1 association in *wpl1*Δ and *elg1*Δ *wpl1*Δ mutants compared to WT specifically at the pericentromeric region (Supplementary Figure S2D), but not at the chromosome arm (Supplementary Figure S2E). Depletion of Pds5 using an auxin-inducible degron system in G1-arrested cells was shown previously to increase genome-wide Scc1 chromatin association two-fold (40). The authors suggested that Pds5 negatively regulates Scc2-mediated cohesin loading throughout the genome. In line with their findings, we observed a two-fold increase in Scc1 chromatin levels at the centromere-distal region in nocodazole-arrested *elg1*Δ *wpl1*Δ *pds5*Δ mutant compared to WT and *elg1*Δ *wpl1*Δ cells (Supplementary Figure S2E). Scc1 chromatin levels at the pericentromeric region in the *elg1*Δ *wpl1*Δ *pds5*Δ were also increased at least two-fold compared to WT, however no further increase was observed compared to *elg1*Δ *wpl1*Δ double mutant (Supplementary Figure S2D). Elevated Scc1 chromatin loading mediated by Scc2 in the absence of Pds5 as assessed by ChIP does not however provide information regarding the fate of the loaded cohesin complexes, which may be targeted by subsequent SUMO-chain-mediated turnover in *PDS5* null cells causing a major reduction of Eco1-mediated Smc3 acetylation levels (Figure 2D and Supplementary Figure S2C).

### Ulp2 protease is essential in *pds5*Δ *elg1*Δ *wpl1*Δ cells and its loss is bypassed by spontaneous suppressor of *ulp2*Δ

How the *elg1*Δ *wpl1*Δ double mutant bypasses the essential role of *PDS5* in counteracting SUMO chains is unclear, but we speculated that the viability of *pds5*Δ *elg1*Δ *wpl1*Δ triple mutant may depend on the presence of the Ulp2 protease as the only source of SUMO-chain-editing activity in yeast cells. To test our hypothesis, we replaced the endogenous promoter of *ULP2* with the galactose-inducible *GAL* promoter, whose transcription can be inhibited by shifting cells from galactose-containing YP Gal to glucose-containing YPD media. Importantly, upon transcriptional *ULP2* shut-off, the *elg1*Δ *wpl1*Δ *pds5*Δ mutant lost viability, whereas *elg1*Δ *wpl1*Δ *PDS5* cells were not affected (Figure 2F). Furthermore, the lethality of *elg1*Δ *wpl1*Δ *pds5*Δ mutant following Ulp2 depletion could be suppressed by a spontaneous suppressor of *ulp2*Δ phenotypes that we identified and denoted as *ulp1-C376* (Figure 2G), described below (Supplementary Figure S3).

Cells lacking Ulp2 exhibit a pleiotropic phenotype that includes temperature-sensitivity and increased sensitivity to hydroxyurea (41). We performed a screen for spontaneous suppressors of *ulp2*Δ-associated HU sensitivity and isolated 5 suppressors that also alleviated the temperature sensitivity (Supplementary Figure S3A). Backcrossing isolated suppressors to the *ulp2*Δ mutant of the opposite mating type, revealed 2^+^:2^−^ segregation of HU sensitivity (Supplementary Figure S3B), which points to a single mutated gene locus responsible for the suppression. Whole genome sequencing (WGS) of the isolated suppressors led to the identification of a single point mutation on chromosome 16 at the *YPL020C* (*ULP1*) gene in all five suppressors, but not in WT or *ulp2*Δ cells (Supplementary Figure S3C). Ulp1 is the second yeast SUMO protease besides Ulp2. Differently from Ulp2, it does not have preference for SUMO chains, is essential for the maturation of conjugatable SUMO from its precursor polypeptide, and is anchored to nuclear pore complexes (NPC) via its N-terminus (41–45). Re-sequencing of the *ULP1* locus validated the WGS results and confirmed a single insertion c.741_742insA denoted as *ulp1-sup* (Supplementary Figure S3D). This insertion results in a frame shift predicted to generate a C-terminally truncated Ulp1 variant p.Val248Serfs*7 lacking the protease domain (Supplementary Figure S3E), a notion not consistent with the essential nature of Ulp1 and its protease domain (Supplementary Figure S3F-G) and different from the isolated *ulp1-sup* (Supplementary Figure S3H-I). Careful analysis of the *ULP1* sequence revealed an alternative transcription/translation start site upstream of the frame shift mutation that might be used in *ulp1-sup* causing the expression of an N-terminally truncated Ulp1 variant (Supplementary Figure S3I) with a new N-terminal extension of 7 amino acids upstream of Lys246 (Supplementary Figure S3J). To confirm that *ulp1-sup* is indeed generating an N-terminally truncated Ulp1 variant able to suppress *ulp2*Δ phenotypes and provides viability as the only source of SUMO protease activity in the cell, we constructed the *ulp1-C376* mutant (Supplementary Figure S3K) by replacing *ulp1*Δ in *ULP2*/*ulp2*Δ *ULP1*/*ulp1*Δ diploid cells with *ulp1* sequence c.[741_742insA; 1_712del] that lacks the N-terminal NPC-targeting region (aa 1-245; first 712 nucleotides of *ULP1* ORF deleted). Tetrad dissection of the resulting *ULP2*/*ulp2*Δ *ULP1*/*ulp1-C376* diploids (Supplementary Figure S3L) revealed that *ulp1-C376* mutant suppressed the HU and temperature sensitivity of *ulp2*Δ cells similar to *ulp1-sup* (Supplementary Figure S3M). Because Ulp1-C376 is expressed at very low levels from the alternative start site compared to wild-type Ulp1 (Supplementary Figure S3I), it does not lead to cell death due to excessive deSUMOylation observed upon expression of N-terminal Ulp1 truncations at higher levels (43, 45).

Taken together, the identified spontaneous suppressor of *ulp2*Δ phenotypes *ulp1-C376* is the N-terminal truncation of Ulp1 that is no longer tethered to the NPC and is able to deSUMOylate Ulp2 substrates in the nucleoplasm providing viability to otherwise lethal *elg1*Δ *wpl1*Δ *pds5*Δ *ulp2* quadruple mutant (Figure 2G). Thus, the viability of *pds5*Δ *elg1*Δ *wpl1*Δ cells depends on Ulp2 ability to counteract SUMO chains and can be bypassed by inducing limited deSUMOylation with the *ulp1-C376* suppressor of *ulp2* phenotypes.

### Loss of Pds5 triggers SUMOylation of cohesin’s kleisin Scc1 that is counteracted by Ulp2

We showed that SUMOylated cohesin subunits are targeted by SUMO chains for turnover in the absence of Ulp2 (Figure 1); that the cohesin’s regulatory subunit Pds5 is no longer essential when SUMO chains cannot form (Figure 2B); and that *pds5*Δ *elg1*Δ *wpl1*Δ cells rely on Ulp2 activity or may survive when N-terminal Ulp1 truncation not tethered to the NPC can perform Ulp2 functions (Figure 2F-G). Next, we aimed to address whether Ulp2 together with Pds5 indeed protects SUMOylated cohesin from SUMO-chain-mediated turnover.

In line with previous observations in a *pds5-1* mutant (11), we find that the lethality due to transcriptional *PDS5* shut-off is suppressed by *slx5*Δ and expression of chainless SUMO *smt3-KRall* variant (Supplementary Figure S4A). Importantly, conditional depletion of Pds5 expressed under the *GAL* promoter allowed us to follow the induction of cohesin SUMOylation and turnover (Figure 3A) by monitoring SUMO modification of cohesin’s kleisin Scc1 and its Slx5/8-targeted proteasome-mediated degradation, previously shown to be enhanced in *pds5-1* (11). Scc1 monoSUMOylation is detected at low levels prior to *PDS5* shut-off (induced by shift to glucose-containing YPD media) in cells expressing ^His^SUMO able to form chains (Figure 3A). Four-eight hours after the shift to YPD, Pds5 levels drop while monoSUMOylated Scc1 species massively accumulate and di/triSUMO-modified Scc1 can be detected. Pds5 depletion is accompanied by reduction in Smc3 lysine K112, K113 acetylation levels and degradation of the unmodified Scc1 (Figure 3A, Input), which is hardly detected 20 hours after *PDS5* shut-off and leads to loss of its SUMOylation. Importantly, expression of the chainless SUMO variant *KRall* results in accumulation of monoSUMOylated Scc1 species even prior to Pds5 depletion (Figure 3B, compare time point 0 in *KRall* versus ^His^SUMO background), suggesting that when SUMO-chain formation is prevented, monoSUMOylated Scc1 species become more stable. Moreover, deletion of *ULP2* in the *KRall* background leads to further accumulation of monoSUMOylated Scc1 species now readily detectable also in the Inputs and accompanied by increase in di/triSUMO-modified Scc1 species (Figure 3B, *KRall ulp2*Δ background). The latter suggests that Ulp2 is able to either cleave off SUMO of mono/multiSUMOylated Scc1 or that physical interaction of Ulp2 with cohesin prevents Scc1 SUMOylation, similar to what has been proposed for Pds5 (11). Because expression of the catalytically inactive *ulp2-C624S* mutant results in similar Scc1 multiSUMOylation levels as observed in *ulp2*Δ mutant (Supplementary Figure S4B), we conclude that Ulp2 is able to cleave off SUMO of mono/multiSUMOylated Scc1. Notably, when SUMO chains are able to form (^His^SUMO expressed), the contribution of Ulp2 to the protection of monoSUMOylated Scc1 species is hardly visible in Ni PD and is manifested by the accumulation of diSUMOylated Scc1 species in *ulp2*Δ cells (Supplementary Figure S4C), in contrast to the *KRall* background (Figure 3B, compare time point 0 in *KRall* versus *KRall ulp2*Δ) where mono/multiSUMOylated Scc1 species are no longer converted to the degradation-prone polySUMOylated species and accumulate. Taken together, these data reveal a role for Ulp2 in guarding mono/multiSUMOylated Scc1 species of cohesin against SUMO-chain-mediated proteasomal turnover that is induced upon Pds5 loss.

**Figure 3.**
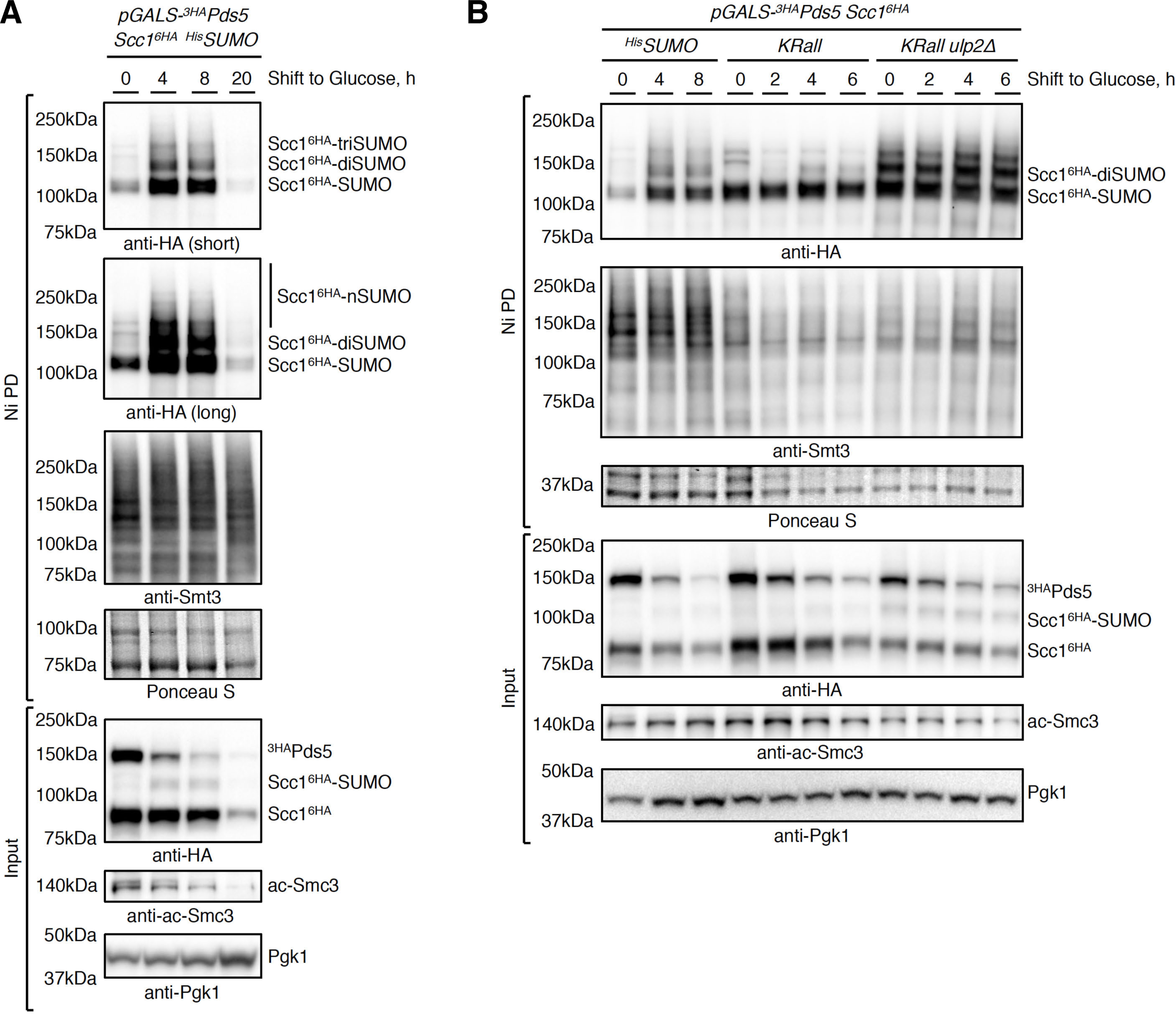
Loss of Pds5 triggers SUMOylation of cohesin’s kleisin Scc1 that is counteracted by Ulp2. (A) Loss of Pds5 triggers SUMOylation of cohesin’s kleisin Scc1 and leads to the reduction of its protein levels as well as of Smc3 lysine K112, K113 acetylation levels. Denaturing Ni-NTA pull-down (Ni PD) was performed to isolate ^His^SUMO conjugates from cells expressing C-terminally 6HA-tagged Scc1 at endogenous levels and N-terminally 3HA-tagged Pds5 under the control of inducible *GAL* promoter. Cells were collected at the indicated time after shift from galactose-to glucose-containing media. Ni PD efficiency was assayed using anti-Smt3 antibody and Ponceau S staining. Pgk1 served as loading control. (B) Expression of a lysine-less SUMO variant *KRall* that cannot form lysine-linked SUMO chains instead of ^His^SUMO results in the accumulation of monoSUMOylated Scc1 species even prior to *PDS5* shut-off. Loss of Ulp2 leads to further increase in Scc1 multiSUMOylation.

### Fusion of Ulp2 to Scc1 supports viability in the absence of Pds5 and protects cohesin from SUMO-chain-mediated turnover

To unambiguously probe whether Ulp2 guards cohesin against SUMO-chain-targeted proteasomal degradation, we next asked if fusion of Ulp2 to the cohesin’s kleisin Scc1 can suppress the lethality of cells lacking Pds5. Previously, fusion of the catalytically active Ulp1 domain to Scc1 (Scc1-UD) was shown to promote loss of cohesin’s SUMO modifications, including monoSUMOylation of Smc3 and Scc1-UD itself (9). Scc1-UD failed to complement the temperature sensitive phenotype of *scc1-73* cells and showed sister chromatid cohesion defects, emphasizing the importance of cohesin SUMOylation for this process. Differently from Scc1-UD that abolishes cohesin SUMOylation, Ulp2 fusion to Scc1 should specifically antagonize its polySUMOylation in the absence of Pds5 leaving mono/multiSUMOylation intact and possibly stabilizing its kleisin.

To this end, we C-terminally fused the Ulp2 fragment (aa 1-734) to the endogenous Scc1 without any linker, providing Scc1-Ulp2 fusion *scc1-ulp2734-6HA* as the only source of cohesin’s kleisin in the cell (Figure 4A). The fused Ulp2 fragment contains N-terminal SIMs required for recognition of SUMO chains (6) and a protease domain, but is lacking its C-terminal 300 amino acids where sequences that mediate its localization to the nucleolus and the inner kinetochore reside (46, 47). Additionally, we established catalytically dead *scc1-ulp2734-C624S-6HA* fusion with Ulp2 active site cysteine mutated (Figure 4A) and *scc1-ulp2734-sim-6HA* fusion having Ulp2 N-terminal SIMs required for robust SUMO-chain-binding mutated (Supplementary Figure S5A).

**Figure 4.**
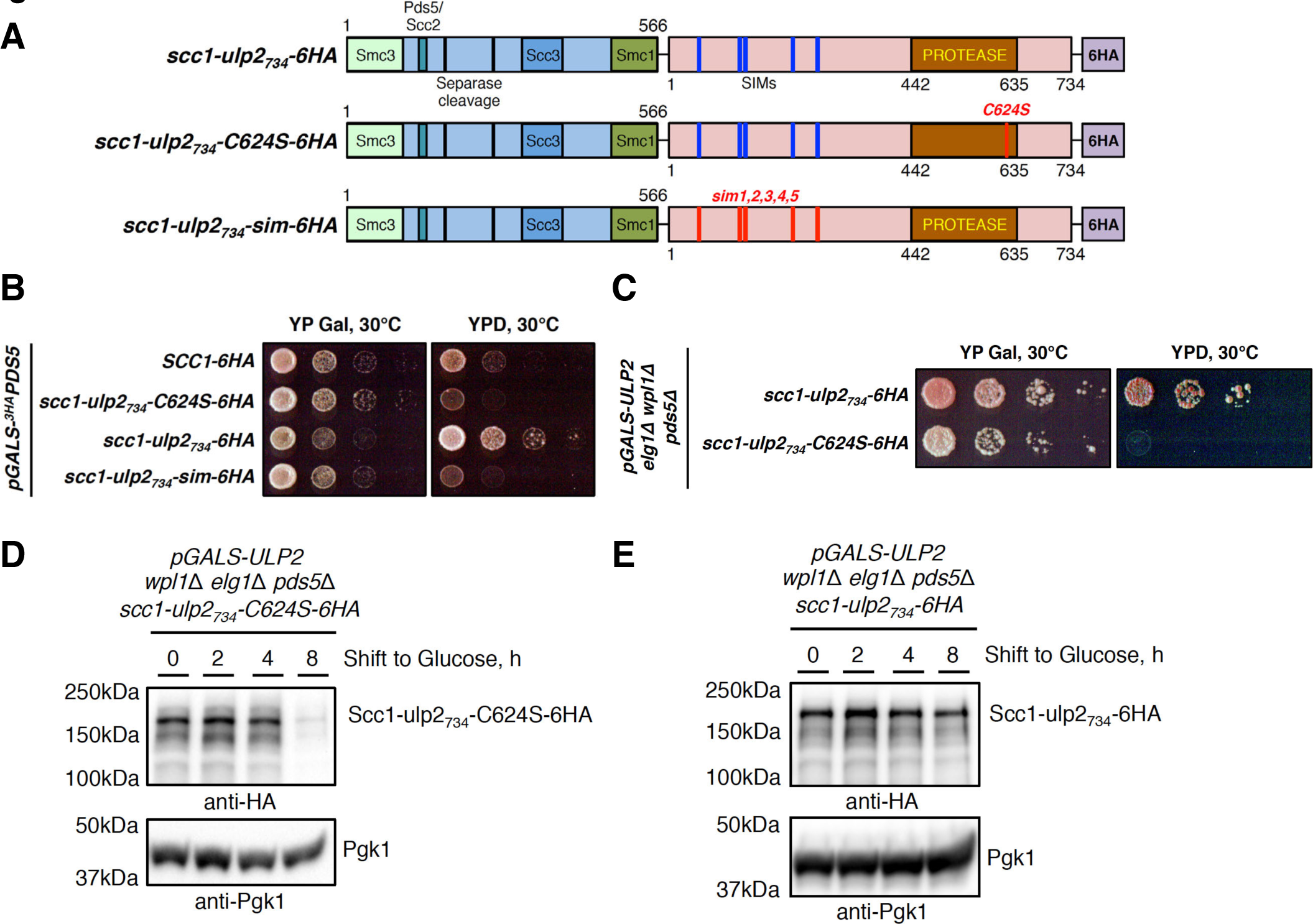
Fusion of Ulp2 to cohesin’s kleisin Scc1 supports viability in the absence of Pds5 and protects cohesin from turnover. (A) Schematic representation of Scc1-Ulp2 fusions used in this study, fusion is the only source of cohesin’s kleisin in the cell. Scc1 depicted with its binding regions to other cohesin subunits and separase cleavage sites. Ulp2 fragment (aa 1-734) is fused C-terminally to endogenous Scc1 without a linker. The N-terminus of Ulp2 harbors 5 SIMs (colored blue) necessary for binding to SUMO chains and is followed by the protease domain; C-terminus (aa 735-1034) is replaced with 6HA tag. Catalytically-active Ulp2 fusion to Scc1 is denoted *scc1-ulp2734-6HA*, catalytically-dead *scc1-ulp2734-C624S-6HA* carries point mutation in active site, *scc1-ulp2734-sim-6HA* has all N-terminal SIMs mutated loosing ability to recognize SUMO chains. (B) Catalytically-active *scc1-ulp2734-6HA* fusion supports viability upon *PDS5* shut-off, whereas *scc1-ulp2734-C624S-6HA* and *scc1-ulp2734-sim-6HA* fusions fail to do so three days post spotting. (C) Catalytically-active *scc1-ulp2734-6HA* fusion supports viability of *pds5*Δ *elg1*Δ *wpl1*Δ cells upon *ULP2* shut-off, whereas catalytically-dead *scc1-ulp2734-C624S-6HA* fusion does not. (D-E) Upon *ULP2* shut-off in *elg1*Δ *wpl1*Δ *pds5*Δ background the protein levels of the catalytically-dead Ulp2 fusion to Scc1 (Scc1-ulp2734-C624S-6HA) decrease (D), whereas catalytically-active Scc1-ulp2734-6HA fusion remains stable (E).

First, we validated that expression of these fusions as the only source of cohesin’s kleisin in WT cells supports viability and that different fusions are expressed at similar levels (Supplementary Figure S5B). Next, we confirmed that expression of the Ulp2 catalytically dead *scc1-ulp2734-C624S-6HA* fusion in the *pds5*Δ *elg1*Δ *wpl1*Δ background is also tolerated (Supplementary Figure S5C).

Importantly, catalytically active *scc1-ulp2734-6HA*, but not catalytically dead *scc1-ulp2734-C624S-6HA* nor *scc1-ulp2734-sim-6HA* fusion defective in binding to SUMO chains, was able to efficiently suppress the lethality of cells upon *PDS5* transcriptional shut-off (Figure 4B). Interestingly, if cells were allowed to grow longer on YPD plates, minor suppression of lethality induced by Pds5 depletion was also observed in cells expressing *scc1-ulp2734-sim-6HA* fusion (Supplementary Figure S5D), suggesting that when fused to Scc1, Ulp2 is able to antagonize polySUMOylation even when SUMO-chain recognition is compromised. In the same line, *scc1-ulp2734-6HA* fusion, but not catalytically dead *scc1-ulp2734-C624S-6HA* provided for viability of *pds5*Δ *elg1*Δ *wpl1*Δ cells upon glucose-induced transcriptional shut-off of *ULP2* (Figure 4C).

Finally, we followed the turnover of the Scc1-Ulp2 fusions in *pds5*Δ *elg1*Δ *wpl1*Δ cells upon *ULP2* shut-off and found that while the catalytically dead fusion is largely degraded 8 hours after Ulp2 depletion (Figure 4D), the catalytically active fusion remains stable (Figure 4E).

Moreover, we followed the SUMOylation status of these fusions upon *ULP2* shut-off by performing Ni PD of ^His^SUMO conjugates able to form SUMO chains (Figure 5A-B). Catalytically active Scc1-Ulp2 fusion remained monoSUMOylated (Figure 5A, Ni PD) and Smc3 lysine K112, K113 acetylation levels did not change significantly upon depletion of Ulp2 (Figure 5A, Input). In contrast, catalytically dead Scc1-Ulp2 fusion became excessively SUMOylated after four hours of Ulp2 depletion (Figure 5B, Ni PD), monoSUMOylated species were converted to slower-migrating polySUMOylated species and finally degraded resulting in the loss of unmodified Scc1-Ulp2 fusion after eight and twenty hours (Figure 5B, Input). Smc3 lysine K112, K113 acetylation levels were also largely reduced (Figure 5B), similar to the situation upon *PDS5* shut-off (Figure 3A). Taken together, these experiments demonstrate that Ulp2 protects cohesin from SUMO-chain-mediated proteasomal turnover in collaboration with Pds5, and that the essential function of budding yeast Pds5 is to counteract SUMO-chain targeting of cohesin, which can be bypassed either by expression of chainless SUMO variant (Figure 2B) or by the Ulp2 fusion to cohesin.

**Figure 5.**
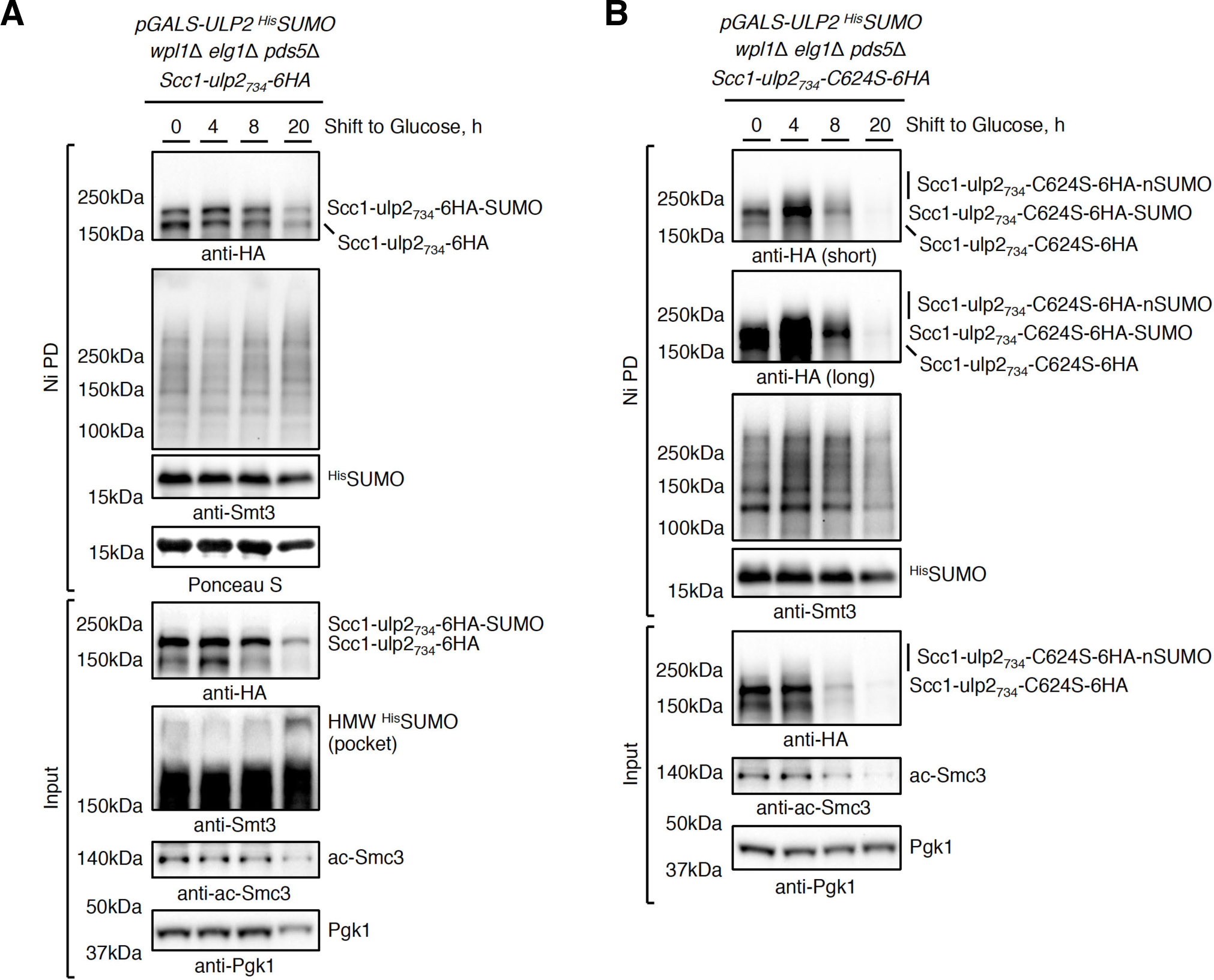
Ulp2 fusion to cohesin’s kleisin Scc1 keeps it in monoSUMOylated state and protects from SUMO-chain-targeted turnover. (A-B) Catalytically-active *scc1-ulp2734-6HA* fusion maintains its protein and monoSUMOylation levels as well as Smc3 lysine K112, K113 acetylation levels upon *ULP2* shut-off in *pds5*Δ *elg1*Δ *wpl1*Δ cells, whereas catalytically-dead *scc1-ulp2734-C624S-6HA* fusion does not. Ni PD was performed to isolate ^His^SUMO conjugates from cells expressing either *scc1-ulp2734-6HA* (A) or *scc1-ulp2734-C624S-6HA* (B) and *ULP2* under the control of inducible *GAL* promoter. Ni PD efficiency was assayed using anti-Smt3 antibody and Ponceau S staining. Pgk1 served as loading control. High molecular weight (HMW) polySUMO conjugates accumulate in Input upon *ULP2* shut-off.

### Condensin is protected by Ulp2 against SUMO-chain-mediated turnover

Condensin subunits are known SUMO substrates (22), however the functional significance of their modification is not clear. Recently, suppressor screening in fission yeast revealed that mutants of condensin’s non-SMC subunits are rescued by impairing the SUMOylation pathway (48). Moreover, deleting *ULP2* was synthetically lethal with the fission yeast temperature sensitive *cut3/smc4* mutant at permissive temperatures for *cut3-477*, while the *ulp2*Δ mutant showed defective chromosome condensation (49).

In our SILAC-based proteomics screen, we identified condensin subunits Smc4 and Brn1 as potential substrates of SUMO-chain-targeted turnover in cells lacking Ulp2 (Figure 1). Furthermore, overexpression of *ULP2* was found to suppress the temperature sensitivity of the *smc2-6* mutant, while *ulp2*Δ cells were defective in enriching condensin on mitotic chromatin, in particular at rDNA (50, 51). We reasoned that similar to cohesin, the condensin complex may be guarded by Ulp2 against SUMO-chain-mediated turnover. We first used genetic analysis to strengthen our hypothesis. While *ULP2* overexpression suppresses *smc2-6* (50) and *smc2-8* (14) alleles, we found that *ulp2*Δ has synergistic growth defect with temperature sensitive *smc2-8* mutant (Figure 6A) at temperatures permissive for the single mutants, similar to fission yeast *cut3/smc4* (49), suggesting that in the absence of Ulp2 the function of condensin is further compromised in *smc2-8* cells. Importantly, the above-mentioned synthetic lethality of *smc2-8 ulp2*Δ cells is rescued by expressing as the single source of SUMO the *smt3-KRall* SUMO variant that cannot form lysine-linked SUMO chains (Figure 6B).

**Figure 6.**
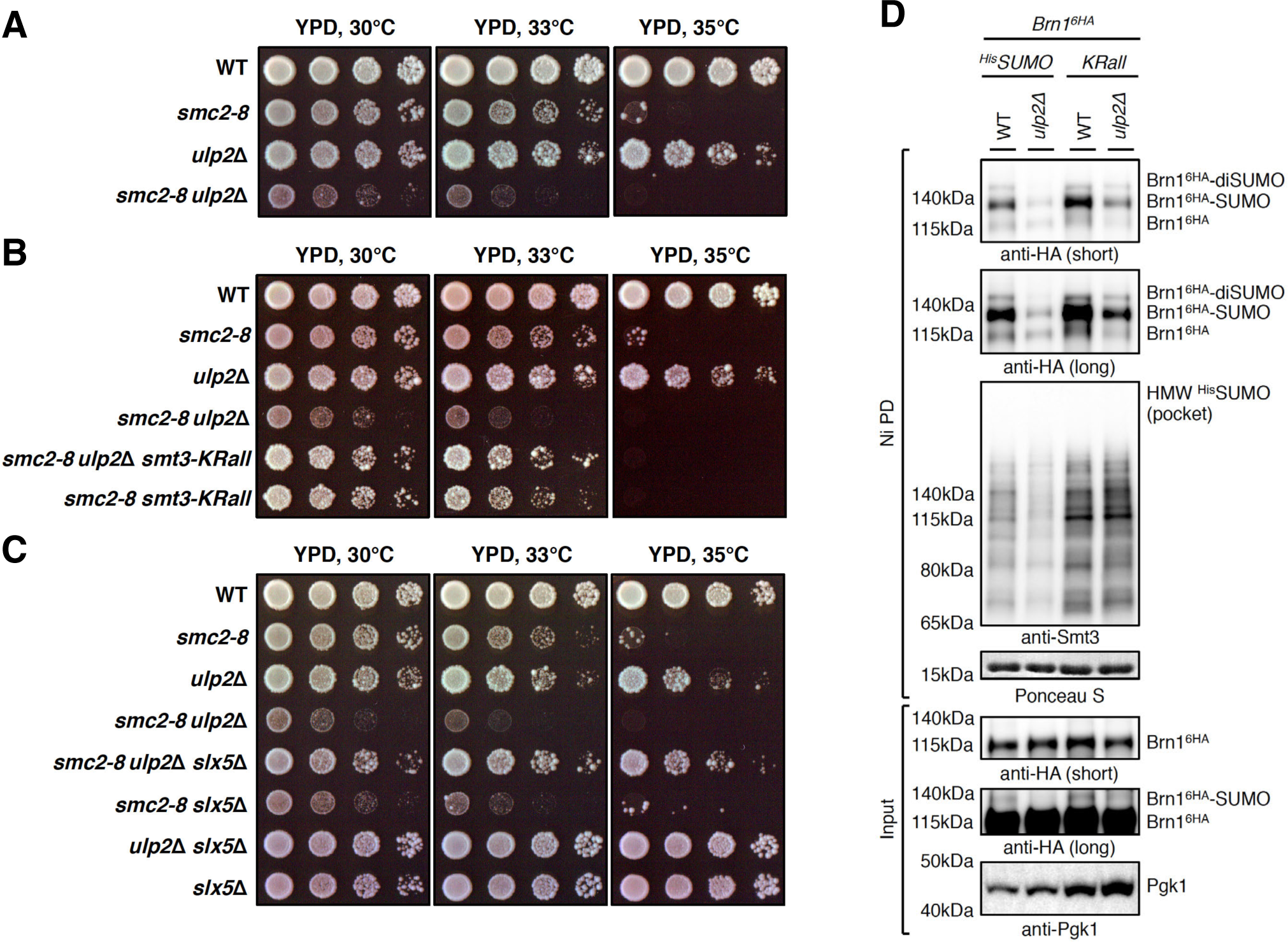
Condensin is protected by Ulp2 against SUMO-chain-mediated turnover. (A) Temperature sensitive *smc2-8* mutant exhibits synthetic sick/lethal genetic interaction with *ulp2*Δ at permissive temperatures for *smc2-8* cells. Spotting of 1:7 serial dilutions on YPD plates at indicated temperatures. (B-C) Synthetic lethality of *smc2-8 ulp2*Δ cells at permissive temperatures for *smc2-8* single mutant is suppressed if a lysine-less SUMO variant (*smt3-KRall*) that cannot form SUMO chains is expressed as the only source of SUMO (B) or if STUbL subunit Slx5 is deleted (C). (D) Levels of monoSUMOylated condensin’s kleisin Brn1 are reduced in the absence of Ulp2 compared to WT, while expression of a lysine-less SUMO variant *KRall* restores them. Ni PD was performed to isolate SUMO conjugates from cells expressing C-terminally 6HA-tagged Brn1. Ni PD efficiency was assayed using anti-Smt3 antibody and Ponceau S staining. Pgk1 served as loading control.

In cohesin, the lethality caused by depletion of the HAWK protein subunit Pds5 is suppressed by the expression of *smt3-KRall* (Figure 2B) and is accompanied by massive SUMOylation of cohesin’s kleisin Scc1 (Figure 3). However, this was not the case when we conditionally depleted the essential HAWK proteins of condensin, Ycg1 and Ycs4, in the *smt3-KRall* background (Supplementary Figure S6A), emphasizing that protection of condensin against SUMO chains is not their essential function. Furthermore, multiSUMOylation of condensin’s kleisin Brn1 tagged C-terminally with 6HA slowly decreased following transcriptional shut-off of either *YCG1* (Supplementary Figure S6B) or *YCS4* (Supplementary Figure S6C), contrary to multiSUMOylation of cohesin’s kleisin Scc1 upon Pds5 depletion (Figure 3A). Nevertheless, the synthetic lethality of *smc2-8 ulp2*Δ cells was suppressed by expressing *smt3-KRall* (Figure 6B) and deleting *SLX5* (Figure 6C and Supplementary Figure S6D). Taken together, these genetic studies support a model in which Ulp2 guards condensin against SUMO-chain-targeted and Slx5/8 STUbL-mediated turnover. Corroborating this, loss of Ulp2 in cells expressing ^His^SUMO able to form chains resulted in the decrease of monoSUMOylated Brn1 species compared to WT cells detected by Ni PD, whereas expression of the chainless SUMO mutant *KRall* not only increased the abundance of monoSUMOylated Brn1 in WT cells, but also suppressed the observed drop in *ulp2*Δ mutant (Figure 6D). However, monoSUMOylated Brn1 species are not particularly prone to proteasomal degradation, as they are not stabilized in the temperature sensitive *cim3-1* proteasome defective mutant (Supplementary Figure S6E) grown at permissive temperature. Overall, the results suggest that in the absence of Ulp2, SUMO chains might target the condensin complex for disassembly and release from chromatin.

### The Smc5/6 complex is protected by Ulp2 against SUMO-chain-mediated turnover

We next studied if the third SMC complex Smc5/6 is similarly regulated. First, using a genetic approach, we found that *ulp2*Δ has synergistic growth defects with the temperature sensitive *smc6-56* mutant (Figure 7A) at temperatures permissive for the single mutants, suggesting that in the absence of Ulp2, the function of the Smc5/6 complex is further compromised, similar to condensin (Figure 6A).

**Figure 7.**
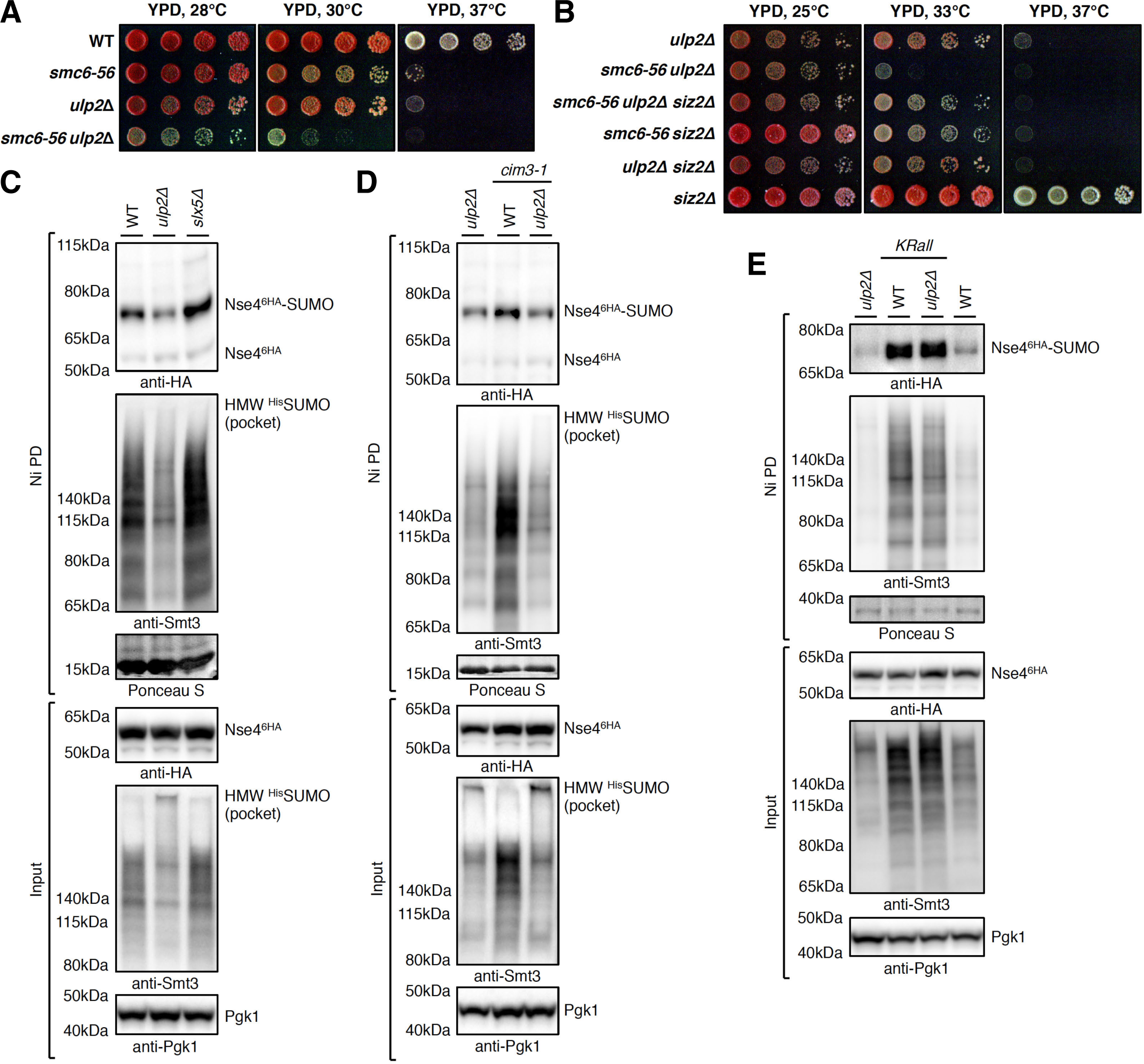
The Smc5/6 complex is protected by Ulp2 against SUMO-chain-mediated turnover. (A) Temperature sensitive *smc6-56* mutant exhibits synthetic sick/lethal genetic interaction with *ulp2*Δ at permissive temperatures for *smc6-56* cells. (B) Synthetic lethality of *smc6-56 ulp2*Δ cells at permissive temperatures for *smc6-56* single mutant is suppressed by deletion of the SUMO ligase Siz2. (C) Levels of monoSUMOylated Smc5/6 complex kleisin Nse4 are reduced in the absence of Ulp2 compared to WT and increased upon Slx5 deletion. Ni PD was performed to isolate ^His^SUMO conjugates from cells expressing C-terminally 6HA-tagged Nse4. Ni PD efficiency was assayed using anti-Smt3 antibody and Ponceau S staining. Pgk1 served as loading control. (D) monoSUMOylated Nse4 species are not degradation-prone as they are not accumulating in the proteasome-defective mutant *cim3-1* background. (E) Expression of a chainless SUMO variant *KRall* leads to the accumulation of monoSUMOylated Nse4 species both in WT and *ulp2*Δ cells.

We then checked if expression of the chainless SUMO variant *smt3-KRall* is able to suppress either *smc6-56* or loss of the essential non-SMC element proteins Nse3 and Nse5. Neither the temperature sensitivity of *nse3-ts-12* and *smc6-56* mutants, nor the lethality upon depletion of Nse5-AID using the auxin-inducible degron system (52) was suppressed by *smt3-KRall* (Supplementary Figure S7A-C). Rather, we observed synthetic sick/lethal genetic interaction between *smc6-56* and *smt3-KRall* at temperatures permissive for the single mutants (Supplementary Figure S7C). Previously, *smc6* mutations were shown to be synthetic sick/lethal with mutation of Sgs1 (53), whereas *sgs1*Δ cells require SUMO-chain formation to survive and die upon expression of *smt3-KRall* (54). Therefore, the synthetic lethality of *smc6-56 smt3-KRall* may be explained by the compromised function that is fulfilled by the Smc5/6 complex in collaboration with Sgs1 (53). The non-SMC element of the Smc5/6 complex Mms21 is a SUMO ligase that is responsible for Smc5/6 complex autoSUMOylation targeting primarily the Smc5 subunit (55). However, the Siz2 SUMO ligase also contributes to Smc5 SUMOylation (56). Because we cannot use *smt3-KRall* to suppress the synthetic lethality of *smc6-56 ulp2*Δ double mutant, we checked if loss of Siz2 is able to do so. Previously, the temperature sensitivity of cohesin’s *pds5-1* mutant was suppressed by *siz2*Δ (11). We found the same to be true also for *smc6-56 ulp2*Δ (Figure 7B) and *smc6-P4 ulp2*Δ (Supplementary Figure S7D) cells. Moreover, the synthetic growth defect of *smc6-P4 ulp2*Δ double mutant is suppressed by deletion of the STUbL subunit Slx5 (Supplementary Figure S7E). In addition, *ulp2*Δ showed synthetic lethal/sick interactions with mutations in Esc2, Sgs1 and Rrm3 (Supplementary Figure S7F-I), previously found to function jointly with Smc5/6 to facilitate resolution of recombination intermediates (Esc2, Sgs1) (57, 58) and replication through difficult-to-replicate regions (Rrm3) (53). Notably, these synthetic sick/lethal interactions could be suppressed by *siz2*Δ (Supplementary Figure S7J-K), *ulp1-C376* (Supplementary Figure S7L) and *slx5*Δ (Supplementary Figure S7M). Taken together, these results suggest that also the Smc5/6 complex is negatively regulated by Siz2-mediated polySUMOylation, which, when not antagonized by Ulp2, is being recognized by the Slx5/8 STUbL and targets the complex for turnover.

In line with genetics, we found that monoSUMOylated species of Nse4, the kleisin of the Smc5/6 complex, accumulate in *slx5*Δ and decrease in abundance in *ulp2*Δ compared to WT cells (Figure 7C). Moreover, similar to condensin’s kleisin Brn1 (Supplementary Figure S6E), monoSUMOylated Nse4 species are not particularly prone to proteasomal degradation as they are not further enriched in *ulp2*Δ *cim3-1* mutant compared to *ulp2*Δ cells (Figure 7D). However, in Ni PD experiments, we find that monoSUMOylated Nse4 strongly accumulates in both WT and *ulp2*Δ when chainless SUMO variant *KRall* is expressed instead of ^His^SUMO (Figure 7E). This indicates that in the absence of Ulp2, SUMO chains might target the Smc5/6 complex for disassembly and release from chromatin, as observed for condensin.

## DISCUSSION

Genetic links between the SUMO protease Ulp2 and components of both condensin (50) and cohesin (14) are known for two decades, yet the role of Ulp2 in the regulation of these SMC complexes has remained elusive. Here, we uncover that Ulp2 acts as a guardian of all three SMC complexes by protecting them from unscheduled SUMO-chain-targeted turnover, thus giving them time to fulfil their functions on chromatin.

Specifically, using unbiased SILAC-based proteomic screen to identify SUMO conjugates that decrease in abundance in the absence of Ulp2 in a SUMO-chain-dependent manner, we found subunits of all SMC complexes (Figure 1). Since *ULP2* overexpression was initially found to suppress phenotypes of *pds5-1* (14), we hypothesized that one important role of Pds5 is to recruit Ulp2 to protect cohesin from polySUMOylation. Strikingly, we showed that Pds5 not only binds Ulp2 (Supplementary Figure S1), but that the essential role of Pds5 is to counteract SUMO-chain formation, as *pds5*Δ lethality is suppressed by expressing a lysine-less SUMO variant (*KRall*) that cannot form SUMO chains (Figure 2B). Moreover, we found that this essential function of Pds5 can by bypassed by combined loss of *ELG1* and *WPL1* (Figure 2C), individual deletions of which suppress cohesion and condensation defects of *pds5-1*, respectively (30). Notably, the viability of *pds5*Δ *elg1*Δ *wpl1*Δ cells relies on Ulp2 (Figure 2F) or could be supported in its absence by expressing the *ulp1-C376* spontaneous suppressor of *ulp2*Δ phenotypes (Figure 2G), which gains ability to deSUMOylate Ulp2 substrates.

We then uncovered that cohesin’s kleisin Scc1 is in fact a Ulp2 substrate, and loss of Pds5 induces its polySUMOylation and subsequent degradation (Figure 3). Finally, we found that fusion of catalytically active Ulp2 to Scc1 suppresses the lethality of cells upon Pds5 loss. Furthermore, Scc1-Ulp2 fusion rescues the lethality of *pds5*Δ *elg1*Δ *wpl1*Δ mutant upon Ulp2 depletion, and prevents polySUMOylation-targeted degradation of cohesin’s kleisin (Figures 4-5). Thus, our data reveal that Ulp2 guards functional mono/multiSUMOylated cohesin pool loaded onto DNA (9, 10) against unscheduled SUMO-chain-mediated turnover, providing the time window for its action until cohesin presence is no longer required, e.g. in mitosis to allow sister chromatid separation.

Interestingly, Ulp2 interacts with and is negatively regulated by the Polo-like kinase Cdc5 in mitosis, with impact on cohesion (59). *CDC5* overexpression causes centromeric cohesion defects that are suppressed by additionally overexpressing *ULP2*. Moreover, *CDC5* overexpression results in Pds5 dissociation from mitotic chromosomes in pre-anaphase cells, whereas co-overexpression of *ULP2* restores normal Pds5 chromosomal association (59). These findings suggest that Cdc5-mediated inactivation of Ulp2 induces SUMO-chain-targeted and Slx5/8 STUbL-mediated proteasomal degradation of the chromatin-bound mono/multiSUMOylated cohesin pool, ensuring its timely removal from chromosomes in collaboration with separase. Thus, yeast Cdc5 kinase promotes sister chromatid separation by phosphorylating Scc1 to enhance separase-mediated cleavage (60), and in parallel, by inducing SUMO-chain-mediated cohesin turnover through inactivation of Ulp2. Supporting this model, overexpression of *ULP2* in separase mutant *esp1-1* results in synthetic sick genetic interaction even at permissive temperatures for *esp1-1* (11).

If the essential function of Pds5 is to help counteract polySUMOylation of cohesin mediated by Ulp2, why is deletion of *PDS5* lethal whereas *ulp2*Δ cells are viable? The presence of Ulp2, however, becomes essential in the *pds5*Δ mutant, the lethality of which is bypassed by *elg1*Δ *wpl1*Δ (Figure 2F). We explain this by the overall structural organization of the SMC complexes (7, 8) and of cohesin in particular (61, 62), and by the promiscuity of the SUMOylation enzymes (3). Specifically, the SUMO conjugating enzyme Ubc9 efficiently targets any accessible lysine able to enter its active site *in vitro* without additional requirements of SUMO ligases. This feature makes the highly unstructured kleisin with many exposed lysines a perfect substrate for SUMOylation, unless it is bound by structured kleisin-associated regulatory subunits. In fact, human Scc1 is SUMOylated at multiple sites and mutation of 15 lysines to arginines reduces, but does not abolish its SUMOylation (21), whereas yeast Scc1 is multiSUMOylated on at least 11 lysines (10). The deletion of *PDS5* has a number of consequences for cohesin. First, Scc2-mediated cohesin DNA loading is strongly stimulated as there is no Pds5 to antagonize it (40) bringing more cohesin complexes in the vicinity of the DNA-bound SUMO ligases. Second, polySUMOylation at multiple acceptor sites on Scc1 is induced, because Pds5 is no longer available to bind Scc1, whereas Scc2 interaction with Scc1 is likely very dynamic as proposed previously (40). In the above-mentioned study, the authors fail to detect an increase in Scc2 association with the genome upon Pds5 depletion and speculate that Scc2 turnover is too rapid for efficient formaldehyde fixation. Since Scc1 is a highly unstructured protein, its exposed lysines can be readily targeted by the rather promiscuous SUMO conjugation machinery, if not shielded by Pds5. Third, loss of Pds5 abolishes recruitment of its interactors, including the SUMO protease Ulp2, to cohesin. Altogether, loss of Pds5 leads to increased Scc2-mediated DNA loading of cohesin with subsequent polySUMOylation and STUbL-mediated proteasomal degradation, exhausting the available pools of the unmodified Scc1. We show that SUMO-chain build-up and kleisin turnover can be prevented if Ulp2 is fused to Scc1 (Figures 4 and 5), providing viability to cells lacking Pds5.

The suppression of the *pds5*Δ mutant lethality by *elg1*Δ *wpl1*Δ, where polySUMOylation of kleisin still takes place and the presence of Ulp2 is required to antagonize it and support viability, indicates that *elg1*Δ *wpl1*Δ associated suppression is achieved by increasing the amounts of cohesin loaded onto DNA rather than by directly antagonizing polySUMOylation. Wpl1 is a cohesin release factor (32), whereas Elg1 unloads PCNA, which is important for recruiting Eco1 to replication forks to promote cohesion during S phase (63). Interestingly, *ECO1* overexpression suppresses *pds5-1* temperature sensitivity (38), probably via Eco1-mediated Smc3 acetylation that antagonizes Wpl1-mediated cohesin release (36, 37). The increase of cohesin DNA-loading provided by *elg1*Δ *wpl1*Δ is likely restricted to specific loci in yeast, as we did not observe pronounced increase in the Eco1-mediated Smc3 lysine K112, K113 acetylation in *elg1*Δ *wpl1*Δ cells compared to WT (Figure 2D and Supplementary Figure S2C). However, this mild increase in cohesin is sufficient to support viability of *pds5Δ elg1*Δ *wpl1*Δ cells. Deletion of Wpl1 in cells blocked in late G1 was previously shown to cause a major increase specifically in peri-centric cohesin, as monitored by Scc1 chromatin binding (40), whereas conditional depletion of Pds5-AID increased Scc2-mediated cohesin loading throughout the genome two-fold. Interestingly, in line with the above-mentioned findings, we observed statistically significant increase of Scc1 association in *wpl1*Δ and *elg1*Δ *wpl1*Δ mutants compared to WT specifically at the pericentromeric region, but not at the chromosome arm in nocodazole-arrested cells (Supplementary Figure S2D-E), which suggests that *elg1*Δ *wpl1*Δ might provide viability to *pds5*Δ cells by supporting pericentromeric cohesion. Moreover, we also observed a two-fold increase in Scc1 loading at the centromere-distal region in nocodazole-arrested *elg1*Δ *wpl1*Δ *pds5*Δ mutant compared to WT and *elg1*Δ *wpl1*Δ cells (Supplementary Figure S2E). Interestingly, increased topological DNA association of cohesin in nocodazole-arrested *pds5-101* cells shifted to restrictive temperature compared to WT cells was reported previously (64), as monitored by the accumulation of monomeric supercoiled DNAs (CMs) in a minichromosome assay. Importantly, however, the authors observed in *pds5-101* cells a marked reduction in the accumulation of DNA-DNA concatemers (CDs) that mediate chromatid cohesion by co-entrapment of sister DNAs inside cohesin rings. Thus, despite having increased Scc2-mediated cohesin DNA loading, loss of Pds5 results in decreased Smc3 acetylation levels (Figure 2D and Supplementary Figure S2C), loss of sister chromatid cohesion as monitored by minichromosome assay (64) and cell lethality. In our work, we demonstrate that the lethality of *pds5*Δ cells is suppressed by expression of a SUMO variant unable to form chains (Figure 2B) or by fusing the SUMO protease Ulp2 that trims SUMO chains to the cohesin’s kleisin Scc1 (Figure 4B-C). The fusion of catalytically active Ulp2 to Scc1 prevents its polySUMOylation and subsequent Scc1 kleisin turnover via proteasomal degradation (Figure 5), providing viability to cells lacking Pds5.

Similar to cohesin, Ulp2 guards the other two SMC complexes, condensin and the Smc5/6 complex, protecting them from SUMO-chain-targeted Slx5/8 STUbL-mediated turnover, as demonstrated by genetic interaction studies and the analysis of their kleisin SUMOylation in relevant mutants of the SUMO/ubiquitin pathway. The mechanism of Ulp2 action is similar, yet it likely results in preventing the disassembly and removal of the complexes from chromatin, rather than massive proteasomal degradation. Accordingly, monoSUMOylated species are not further enriched in the proteasome defective *cim3-1* mutant, but are nevertheless stabilized upon expression of the chainless SUMO variant *KRall*. Chromatin extraction of polySUMOylated and subsequently polyubiquitylated SMC complexes is likely mediated by the action of the Cdc48/p97 segregase (65, 66), which was previously shown to mobilize cohesin and condensin from chromatin (67, 68). Despite this difference, the outcome of Ulp2 loss for all SMC complexes is the same in that their turnover on chromatin is increased. Another difference is that depletion of condensin’s HAWKs or the KITEs of the Smc5/6 complex is not suppressed by expressing the *KRall* SUMO mutant, suggesting that it is not their essential function to counteract kleisin polySUMOylation, contrary to Pds5. We note that Pds5 is likely a special case, as it is not stimulating the ATPase activity required for the DNA-loading of cohesin, in contrast to the Scc2 HAWK, with which Pds5 competes and then replaces to stabilize cohesin complex in the DNA-bound state (40).

In conclusion, here we uncovered that the SUMO protease Ulp2 acts on all three SMC complexes to protect them from unscheduled SUMO-chain-targeted turnover, giving them time to perform their essential functions on chromatin. Since Ulp2 discovery two decades ago (41), multiple phenotypes have been associated with its loss, including genome instability, sister chromatid cohesion and condensation defects. Our finding helps to explain many of them and highlights a new layer of the SMC complex regulation, through which their chromatin abundance is instructed.

## SUPPLEMENTARY DATA

Supplementary Data are available at NAR online.

## ACKNOWLEDGEMENTS

We thank Eurofins Genomics for the whole genome sequencing of yeast strains, mapping the sequencing reads and variant analysis, M. Foiani, S. Jentsch, and K. Shirahige for sharing reagents, A. Cattaneo and A. Bachi for mass spectrometry analysis, L. Fendillo for help with generating Scc1-Ulp2 fusions, R. Kawasumi and all lab members for discussions.

