## Supplemental information for "SMC Complexes Are Guarded by the SUMO Protease Ulp2 Against SUMO-Chain-Mediated Turnover"

**Supplementary Figures S1-S7**

### Figure S1

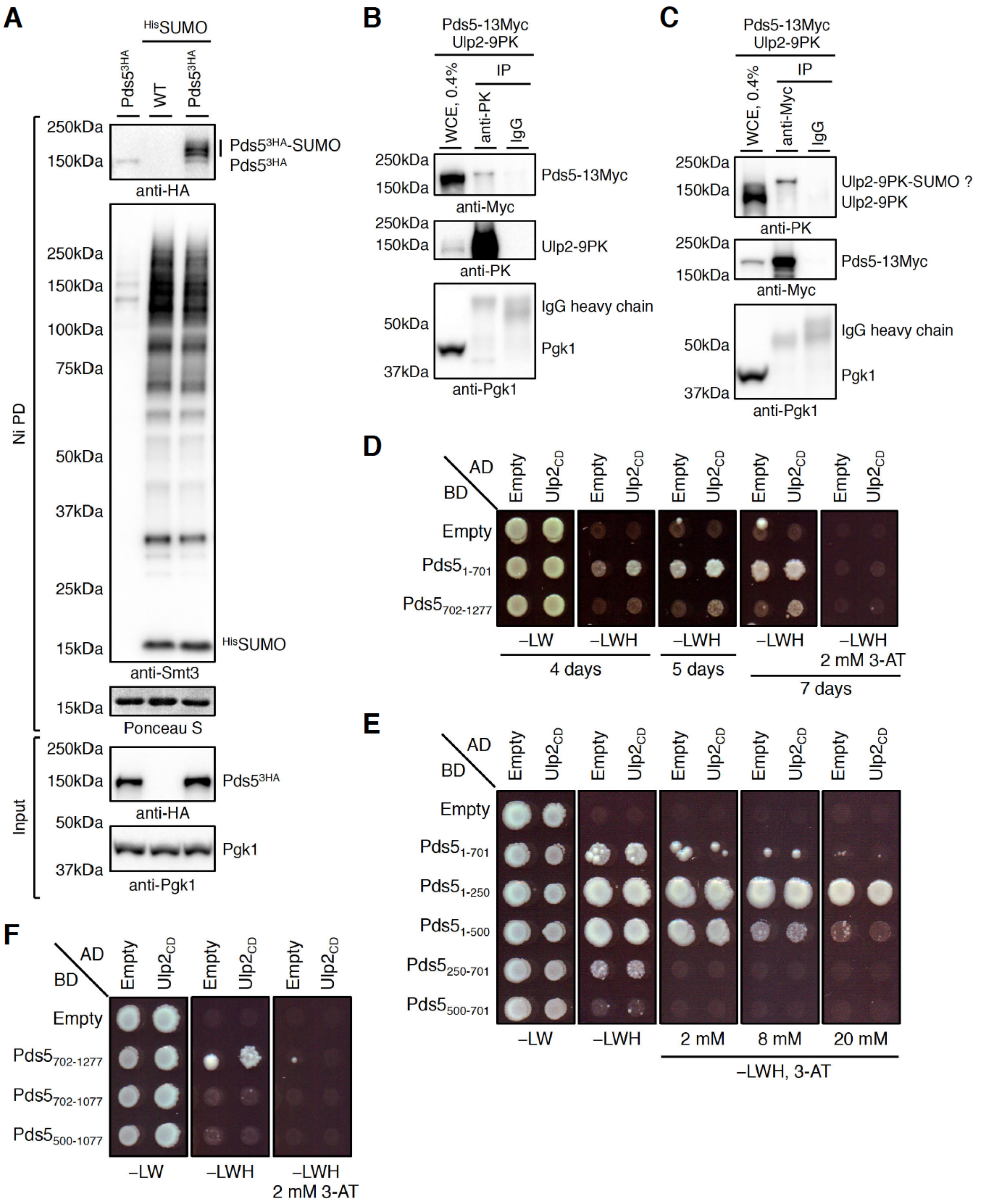
A
HisSUMO
Pds5 3HA
WT
Pds5 3HA
250kDa
150kDa
Pds5 3HA-SUMO
Pds5 3HA
anti-HA
250kDa
150kDa
100kDa
75kDa
50kDa
37kDa
25kDa
15kDa
HisSUMO
anti-Smt3
15kDa
Ponceau S
Ni PD
250kDa
150kDa
Pds5 3HA
anti-HA
50kDa
37kDa
Pgk1
anti-Pgk1
Input
B
Pds5-13Myc
Ulp2-9PK
WCE, 0.4%
IP
anti-PK
IgG
250kDa
150kDa
Pds5-13Myc
anti-Myc
250kDa
150kDa
Ulp2-9PK
anti-PK
50kDa
37kDa
IgG heavy chain
Pgk1
anti-Pgk1
C
Pds5-13Myc
Ulp2-9PK
WCE, 0.4%
IP
anti-Myc
IgG
250kDa
150kDa
Ulp2-9PK-SUMO ?
Ulp2-9PK
anti-PK
250kDa
150kDa
Pds5-13Myc
anti-Myc
50kDa
37kDa
IgG heavy chain
Pgk1
anti-Pgk1
D
AD
BD
Empty
Ulp2CD
Empty
Pds5 1-701
Pds5 702-1277
–LW
–LWH
–LWH
–LWH
–LWH 2 mM 3-AT
4 days
5 days
7 days
E
AD
BD
Empty
Ulp2CD
Empty
Pds5 1-701
Pds5 1-250
Pds5 1-500
Pds5 250-701
Pds5 500-701
–LW
–LWH
2 mM
8 mM
20 mM
–LWH, 3-AT
F
AD
BD
Empty
Ulp2CD
Empty
Pds5 702-1277
Pds5 702-1077
Pds5 500-1077
–LW
–LWH
–LWH 2 mM 3-AT

**Supplementary Figure 1. SUMO protease Ulp2 interacts with cohesin's regulatory subunit Pds5 that undergoes SUMOylation (related to Figure 1)**

(A) Pds5 is SUMOylated. Ni PD was performed to isolate $^{His}$SUMO conjugates from cells expressing C-terminally 3HA-tagged Pds5. Ni PD efficiency was assayed using anti-Smt3 antibody and Ponceau S staining (nonspecifically pulled-down protein of ≈15 kDa visualized). Pgk1 served as loading control.

(B-C) Ulp2 interacts with Pds5 in the co-immunoprecipitation (Co-IP) studies. Whole cell extracts (WCE) were prepared from cells expressing C-terminally 9PK-tagged Ulp2 and 13Myc-tagged Pds5 using grinding in liquid nitrogen followed by solubilization in the lysis buffer and IP. Interaction is observed when tagged proteins are immunoprecipitated either with anti-PK or anti-Myc antibodies, but not with mouse IgG. Co-immunoprecipitated proteins have lower electrophoretic mobility suggesting that they are modified with SUMO.

(D) N-terminal and C-terminal fragments of Pds5 interact with Ulp2 (catalytically-dead variant Ulp2-C624S, $Ulp2_{CD}$, has stronger interaction with its substrates compared to the wild-type protein) in the Y2H system. 2mM 3-amino-triazole (3-AT) was added to selection plates to reduce auto-activation of *HIS3* reporter gene.

(E) N-terminal fragment of Pds5 (amino acids 1-250) shows high auto-activation of *HIS3* reporter gene in the Y2H system even at high concentrations of 3-AT.

(F) C-terminal amino acids 1078-1277 of Pds5 are required for the interaction with Ulp2 in the Y2H system.

Figure S2

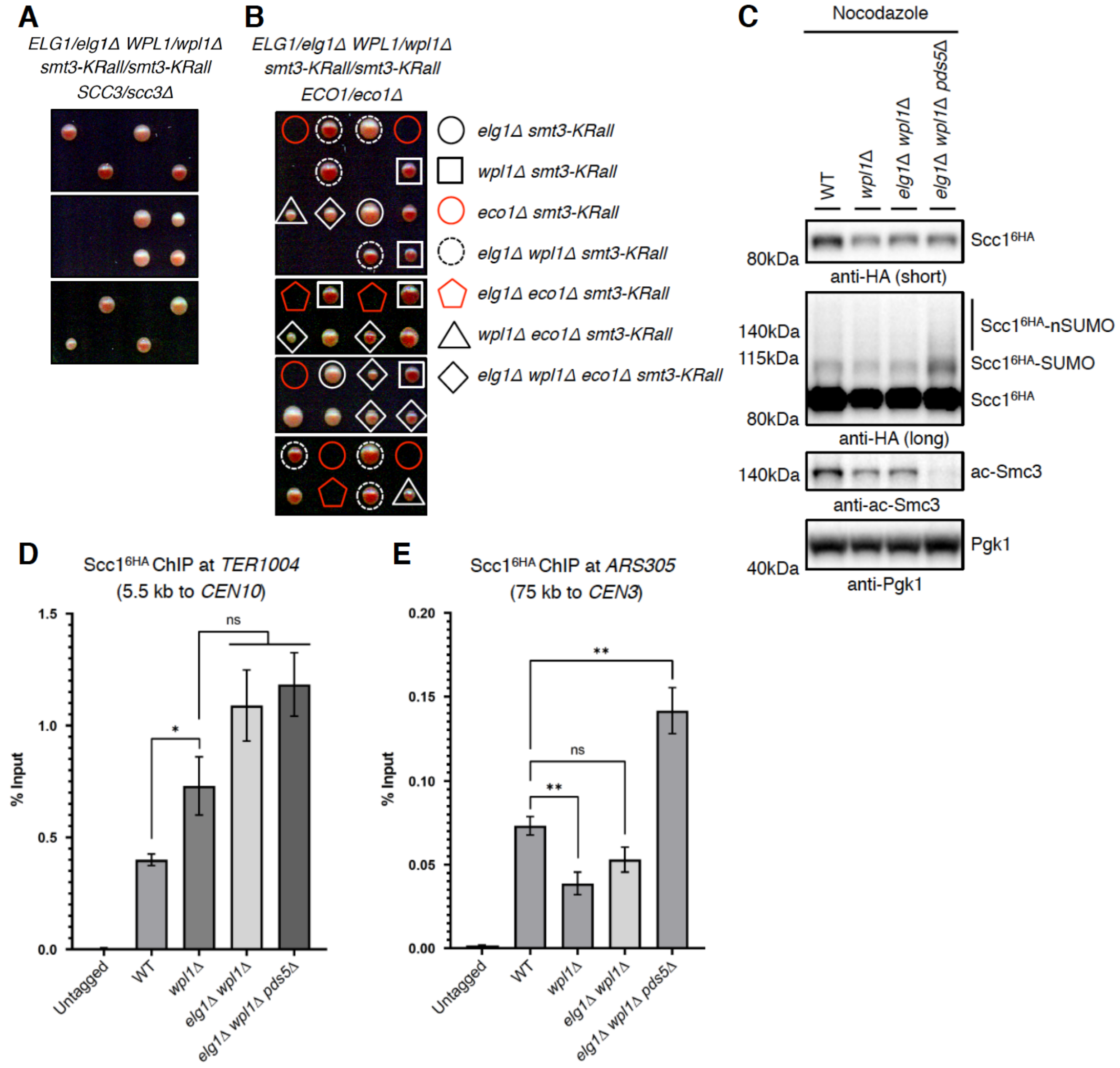


**Supplementary Figure 2. Scc1 loading at the pericentromeric region *TER1004* is increased, but is largely not affected at the centromere-distal region *ARS305* in the *elg1Δ wpl1Δ* cells (related to Figure 2)**

(A) Tetrad dissection analysis of *ELG1/elg1Δ WPL1/wpl1Δ smt3-KRall/smt3-KRall SCC3/scc3Δ* mutant revealed no bypass of *scc3Δ* associated lethality.

(B) Tetrad dissection analysis of *ELG1/elg1Δ WPL1/wpl1Δ smt3-KRall/smt3-KRall ECO1/eco1Δ* mutant. The lethality of *eco1Δ* cells is suppressed only in the absence of *WPL1*.

(C) Smc3 lysine K112, K113 acetylation is reduced in the nocodazole-arrested *elg1*Δ *wpl1*Δ *pds5*Δ cells, accompanied by increase in the Scc1 SUMOylation levels.

(D-E) Binding of $Scc1^{6HA}$ onto chromatin at the pericentromeric region *TER1004* of *CEN10* (D) and the centromere-distal region *ARS305* (E) analyzed in nocodazole-arrested wild type (WT), *wpl1*Δ, *elg1*Δ *wpl1*Δ, and *elg1*Δ *wpl1*Δ *pds5*Δ cells by ChIP-qPCR assay. Untagged Scc1 wild type strain was used as control. Each ChIP experiment was repeated at least three times, and each real-time PCR was performed in triplicates. Statistical analysis was performed using Student's unpaired *t*-test. The mean values ± standard error of mean (SEM) are plotted.

### Figure S3

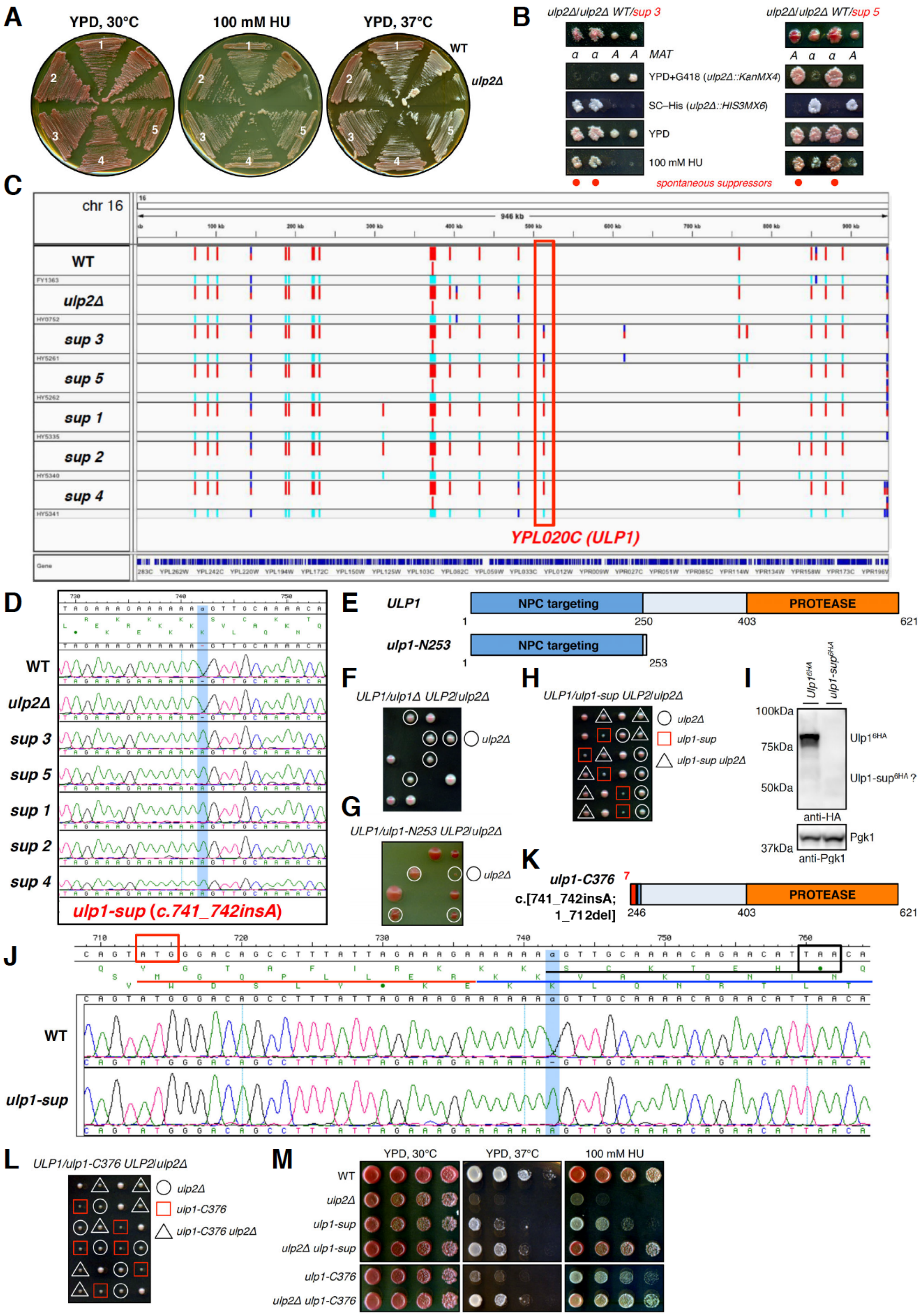
A
YPD, 30°C
100 mM HU
YPD, 37°C
WT
ulp2Δ
B
ulp2Δ/ulp2Δ WT/sup 3
ulp2Δ/ulp2Δ WT/sup 5
MAT
YPD+G418 (ulp2Δ::KanMX4)
SC–His (ulp2Δ::HIS3MX6)
YPD
100 mM HU
spontaneous suppressors
C
chr 16
WT
ulp2Δ
sup 3
sup 5
sup 1
sup 2
sup 4
YPL020C (ULP1)
D
WT
ulp2Δ
sup 3
sup 5
sup 1
sup 2
sup 4
ulp1-sup (c.741_742insA)
E
ULP1
NPC targeting
PROTEASE
ulp1-N253
F
ULP1/ulp1Δ ULP2/ulp2Δ
ulp2Δ
G
ULP1/ulp1-N253 ULP2/ulp2Δ
H
ULP1/ulp1-sup ULP2/ulp2Δ
ulp1-sup
ulp1-sup ulp2Δ
I
100kDa
75kDa
50kDa
37kDa
Ulp1-sup6HA ?
anti-HA
Pgk1
anti-Pgk1
K
ulp1-C376
c.[741_742insA; 1_712del]
J
WT
ulp1-sup
L
ULP1/ulp1-C376 ULP2/ulp2Δ
ulp1-C376
ulp1-C376 ulp2Δ
M
ulp2Δ ulp1-sup
ulp2Δ ulp1-C376

**Supplementary Figure 3. Identification of the spontaneous suppressor of *ulp2*Δ associated phenotypes (related to Figure 2)**

(A-B) Plating of *ulp2*Δ cells on YPD plates containing 100 mM HU has led to the isolation of five spontaneous suppressors able to suppress not only HU, but also the temperature sensitivity (A) and sporulation defects (B) of *ulp2*Δ mutant. Backcrossing of the isolated suppressors to *ulp2*Δ mutant of opposite mating type followed by sporulation and tetrad dissection (B) revealed $2^{+}:2^{-}$ segregation regarding sensitivity to HU that points to a single mutated gene locus responsible for suppression.

(C) View of the indels mapped on chromosome 16 following the whole genome sequencing of WT (W303 background), *ulp2*Δ cells and isolated spontaneous suppressors (*sup 1-5*) of *ulp2*Δ phenotypes. Genome of strain S288C was used as reference for mapping and variant analysis. Point mutation was identified in gene *YPL020C* (*ULP1*) in all suppressors, but not in WT or *ulp2*Δ cells.

(D) Resequencing of *ULP1* gene confirmed single nucleotide insertion c.741_742insA annotated as *ulp1-sup* in all isolated suppressors of *ulp2*Δ phenotypes.

(E) Schematic representation of Ulp1 with its SUMO protease domain and the N-terminal region required for tethering to the nuclear pore complex (NPC). Identified insertion leads to the frame shift and is predicted to result in C-terminally truncated Ulp1 variant p.Val248Serfs*7 (J) lacking protease domain. *ulp1-N253* mutant generated with a stop codon introduced after Ile253.

(F-H) Tetrad dissection analysis of *ULP1*/*ulp1*Δ *ULP2*/*ulp2*Δ (F) and *ULP1*/*ulp1-N253* *ULP2*/*ulp2*Δ (G) cells revealed lethality of spores carrying *ulp1* mutants, whereas dissection of *ULP1*/*ulp1-sup* *ULP2*/*ulp2*Δ diploids with *URA3* marker introduced downstream of *ulp1-sup* resulted in 4 viable spores (H).

(I) *ulp1-sup* mutant is expressed at very low levels. *ULP1* and *ulp1-sup* were C-terminally tagged with 6HA and their expression compared by western blotting. Faint signal detected at around 50 kDa for Ulp1-sup$^{6HA}$ compared to Ulp1$^{6HA}$ at expected 80 kDa.

(J) View of the *ULP1* gene with c.741_742insA mutation highlighted in *ulp1-sup*. Black box indicates predicted stop codon of expected p.Val248Serfs*7 Ulp1 variant with a new C-terminal extension (underlined black); red box indicates potential alternative transcription/translation start site upstream of the frame shift mutation resulting in the expression of an N-terminally truncated

Ulp1 variant with a new N-terminal extension of 7 amino acids (underlined red) upstream of Lys246.

(K) Schematic representation of the generated *ulp1-C376* mutant c.[741_742insA; 1_712del] that lacks N-terminal NPC-targeting region (aa 1-245; first 712 nucleotides of *ULP1* ORF deleted), harbors N-terminal extension of 7 amino acids (red) upstream of Lys246 as in *ulp1-sup* and is expressed from endogenous *ULP1* promoter.

(L) Tetrad dissection of *ULP1*/*ulp1-C376 ULP2*/*ulp2*Δ diploids with *URA3* marker introduced downstream of *ulp1-C376* resulted in 4 viable spores, as in (H) for *ulp1-sup*.

(M) Generated *ulp1-C376* mutant suppresses *ulp2*Δ phenotypes similar to *ulp1-sup* validating the spontaneous suppressor mutation. Spotting of 1:7 serial dilutions on YPD plates at indicated temperatures and HU concentration.

### Figure S4

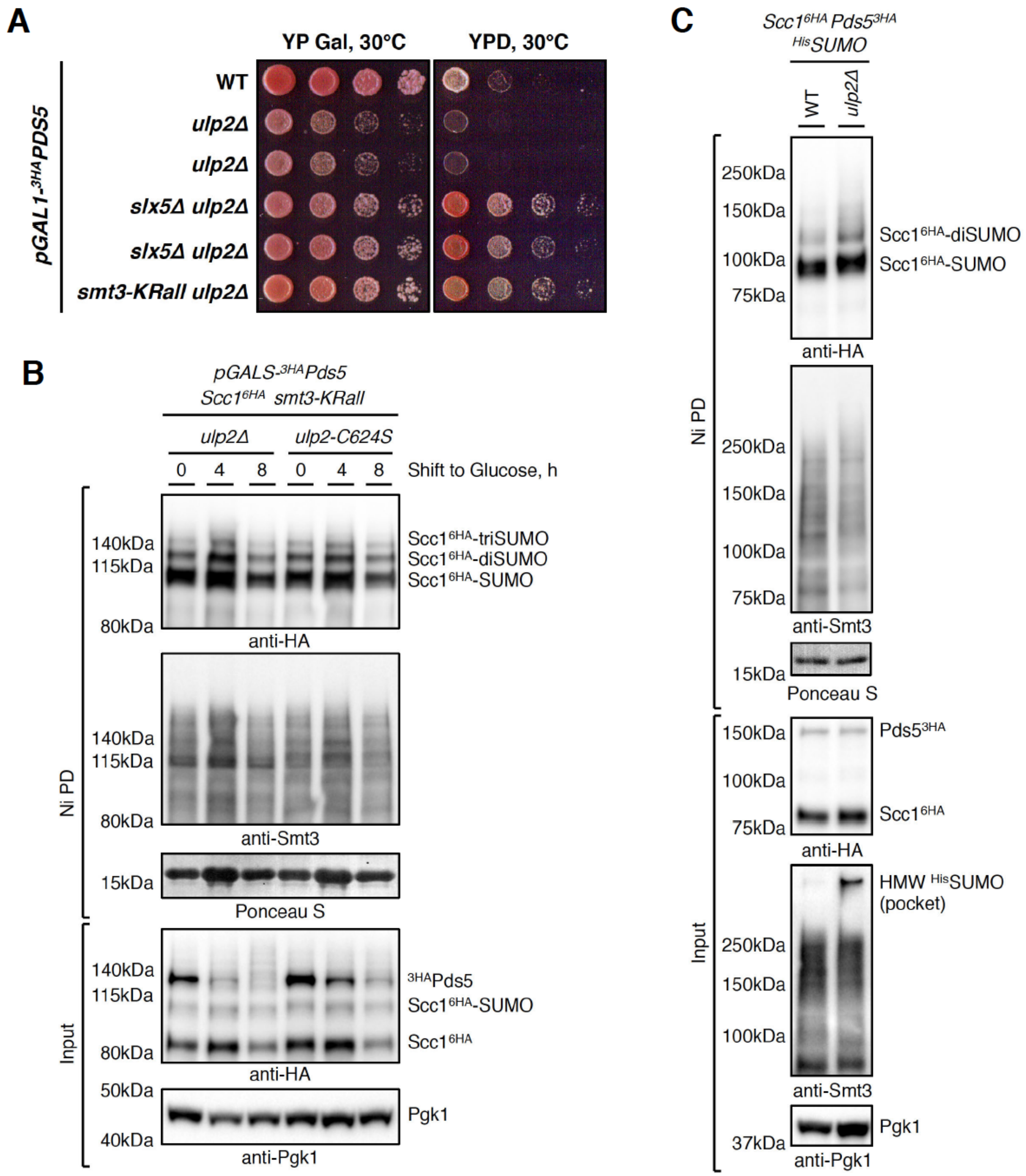


**Supplementary Figure 4. Ulp2 is able to counteract multiSUMOylation of Scc1 (related to Figure 3)**

(A) Upon transcriptional shut-off of *PDS5* expressed from inducible *GAL* promoter WT and *ulp2*Δ cells die, which does not happen in the absence of STUbL subunit Slx5 or when SUMO chains cannot form in *smt3-KRall* mutant.

(B) Expression of the catalytic dead *ulp2-C624S* mutant results in a similar increase of Scc1 multiSUMOylation as observed in *ulp2*Δ cells upon *PDS5* shut-off. Ni PD was performed to isolate chainless SUMO conjugates (*smt3-KRall*) from *ulp2*Δ or *ulp2-C624S* cells with N-terminally 3HA-tagged Pds5 under the control of inducible *GAL* promoter and expressing C-terminally 6HA-tagged Scc1. Ni PD efficiency was assayed using anti-Smt3 antibody and Ponceau S staining (nonspecifically pulled-down protein of ≈15 kDa visualized). Pgk1 served as loading control.

(C) If SUMO is able to form chains, loss of Ulp2 does not lead to pronounced increase in multiSUMOylated Scc1 species compared to WT cells, on the contrary to the situation when chainless SUMO mutant *KRall* is expressed instead (Figure 3B, compare $Scc1^{6HA}$-diSUMO levels in *KRall* versus *KRall ulp2*Δ at 0 h time point). multiSUMOylated Scc1 species are converted to degradation-prone polySUMOylated Scc1 species in *ulp2*Δ cells, detected as up-shifted smear following $^{His}$SUMO Ni PD.

**Figure S5**

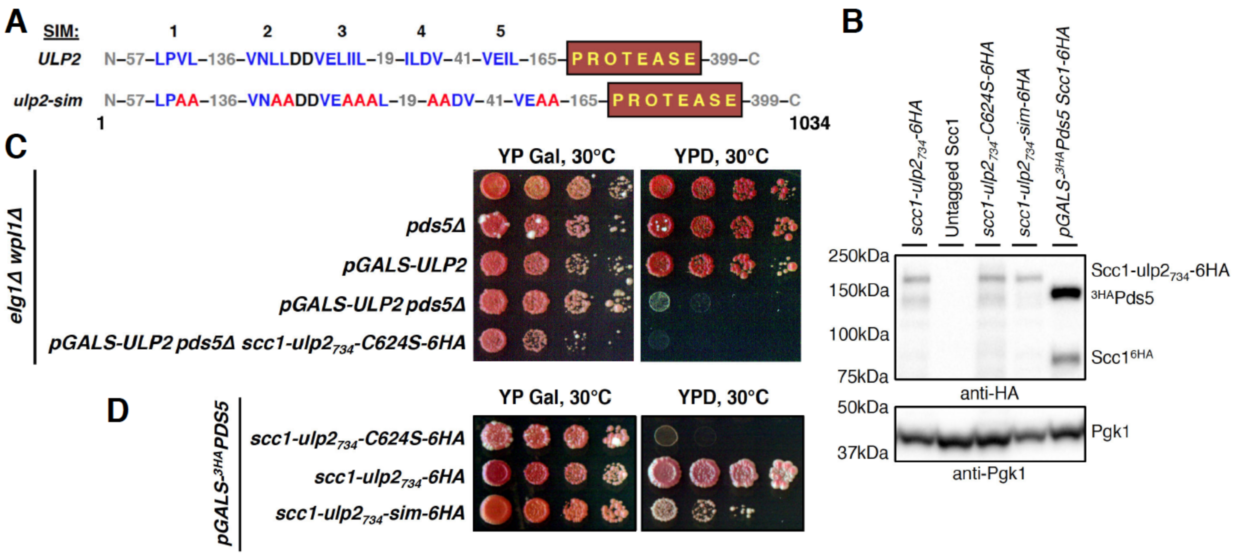


**Supplementary Figure 5. Scc1-Ulp2 fusion protects cohesin from SUMO-chain-mediated turnover upon *PDS5* shut-off or *ULP2* loss in *elg1*Δ *wpl1*Δ *pds5*Δ background (related to Figures 4 and 5)**

(A) Schematic representation of Ulp2 with the protease domain, N-terminal SIMs (colored blue), and introduced point mutations (colored red) that disrupt Ulp2 SIMs required for binding to SUMO chains.

(B) Catalytically-active Ulp2 fusion to Scc1 *scc1-ulp2*$_{734}$*-6HA*, catalytically-dead *scc1-ulp2*$_{734}$*-C624S-6HA,* and *scc1-ulp2*$_{734}$*-sim-6HA* fusion unable to recognize SUMO chains are all expressed at similar levels with endogenous 6HA-tagged *SCC1*, and are the only source of cohesin's kleisin in the cell.

(C) Cells expressing catalytically-dead Ulp2 fusion to Scc1 *scc1-ulp2*$_{734}$*-C624S-6HA* in *elg1*Δ *wpl1*Δ *pds5*Δ background are viable, but are slightly slow-growth compared to cells expressing *SCC1* and similarly do not suppress lethality upon *ULP2* shut-off. Spotting of 1:7 serial dilutions on YP Gal or YPD plates, images taken after 3 days.

(D) Catalytically-active Ulp2 fusion to Scc1 *scc1-ulp2*$_{734}$*-6HA*, but not catalytically-dead *scc1-ulp2*$_{734}$*-C624S-6HA* rescues lethality upon *PDS5* shut-off, whereas *scc1-ulp2*$_{734}$*-sim-6HA* fusion unable to recognize SUMO chains shows weak suppression. Images were taken after 6 days of incubation on YPD plates.

Figure S6

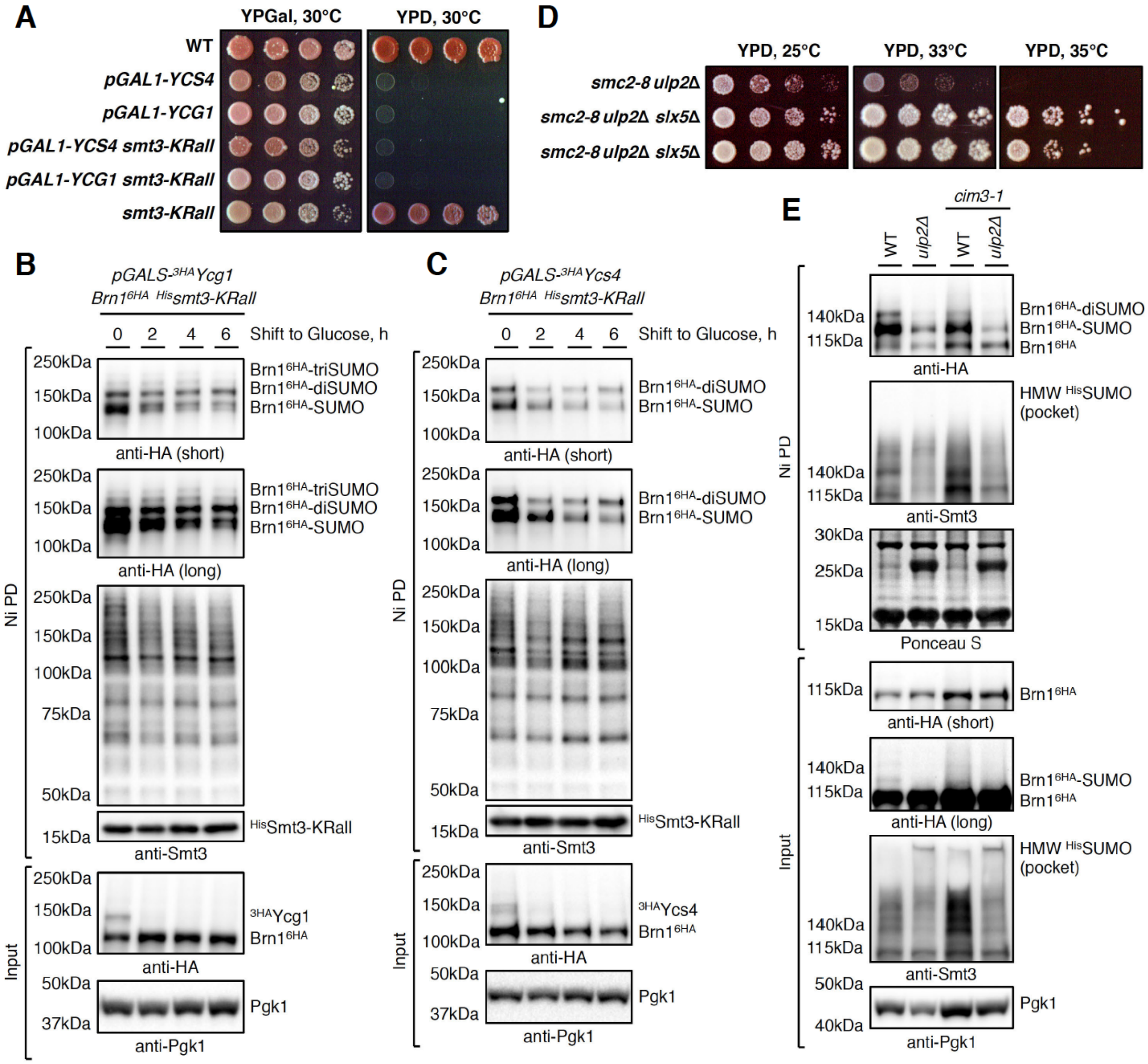


**Supplementary Figure 6. Loss of condensin's HAWK proteins Ycg1 and Ycs4 does not induce kleisin Brn1 SUMOylation and is not bypassed by *smt3-KRall* mutant (related to Figure 6)**

(A) Lethality of cells upon *YCG1* and *YCS4* shut-off is not suppressed by the expression of *smt3-KRall* chainless SUMO mutant. Spotting of 1:7 serial dilutions on YP Gal or YPD plates.

(B-C) Loss of Ycg1 and Ycs4 upon transcriptional shut-off does not induce, but gradually reduces multiSUMOylation of condensin's kleisin Brn1. Ni PD was performed to isolate SUMO conjugates from *smt3-KRall* mutant cells expressing endogenous C-terminally 6HA-tagged Brn1 and N-terminally 3HA-tagged Ycg1 (B) or Ycs4 (C) under the control of inducible *GAL* promoter. Cells were collected at the indicated time after shift from galactose- to glucose-containing media. Ni PD efficiency was assayed using anti-Smt3 antibody. Pgk1 served as loading control.

(D) Synthetic lethality of *smc2-8 ulp2*Δ cells at permissive temperatures for *smc2-8* single mutant is suppressed by Slx5 STUbL subunit deletion. *smc2-8 ulp2*Δ *slx5*Δ triple mutants isolated from different tetrad dissections spotted at indicated temperatures.

(E) monoSUMOylated Brn1 species are not prone to proteasomal degradation as they are not accumulating in the temperature sensitive proteasome-defective mutant *cim3-1* background. Ni PD of $^{His}$SUMO conjugates from WT, *ulp2*Δ, *cim3-1* and *cim3-1 ulp2*Δ cells expressing C-terminally 6HA-tagged Brn1. Cells were grown at the permissive temperature of 30°C for *cim3-1* mutant.

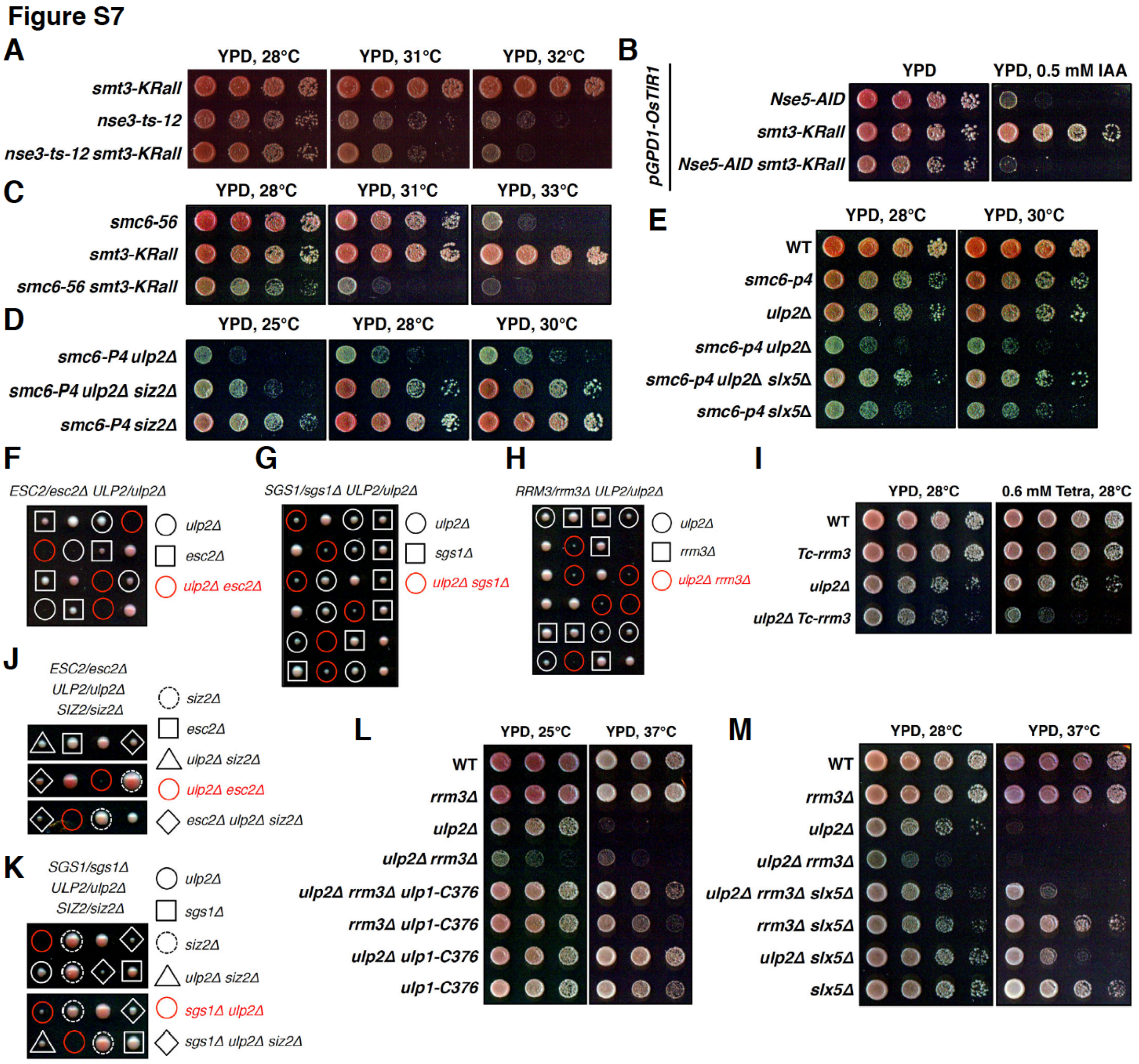


**Supplementary Figure 7. Genetic interaction studies supporting protection of the Smc5/6 complex by Ulp2 protease against SUMO-chain-mediated turnover (related to Figure 7)**

(A) Lethality of temperature-sensitive *nse3-ts-12* mutant is not suppressed by the expression of *smt3-KRall* chainless SUMO mutant. Spotting of 1:7 serial dilutions on YPD plates at indicated temperatures.

(B) Lethality due to depletion of *Nse5-AID* using the auxin-inducible degron system is not suppressed by the expression of *smt3-KRall* mutant. Cells with indicated genotypes expressing *OsTIR1* spotted on YPD plates with or without 0.5 mM IAA.

(C) Lethality of temperature-sensitive *smc6-56* mutant is not suppressed by the expression of *smt3-KRall*, rather *smc6-56 smt3-KRall* double mutant is synthetic lethal at temperature permissive for *smc6-56* single mutant.

(D) Synthetic sick phenotype of *smc6-p4 ulp2*Δ double mutant is suppressed by deleting SUMO ligase Siz2.

(E) Synthetic sick phenotype of *smc6-P4 ulp2*Δ and *smc6-P4 slx5*Δ double mutants is suppressed in the triple mutant.

(F-H) Tetrad dissection analysis of *ESC2/esc2*Δ *ULP2/ulp2*Δ (F), *SGS1/sgs1*Δ *ULP2/ulp2*Δ (G), and *RRM3/rrm3*Δ *ULP2/ulp2*Δ (H) diploids revealed synthetic sick/lethal interactions between *ulp2*Δ and indicated mutants that also exhibit synthetic lethal interactions with *smc6* mutant.

(I) Synthetic sickness of *rrm3*Δ *ulp2*Δ double mutant (H) is confirmed by Rrm3 conditional depletion in *ulp2*Δ cells using *Tc-rrm3* allele that carries a tetracycline-dependent translational repressor. Cells with indicated genotypes spotted on YPD plates with or without 0.6 mM tetracycline.

(J-K) Tetrad dissection analysis of the indicated triple mutants revealed that synthetic sickness/lethality of *esc2*Δ *ulp2*Δ (J) and *sgs1*Δ *ulp2*Δ (K) double mutants is suppressed by deletion of Siz2 SUMO ligase.

(L-M) Synthetic sickness of *rrm3*Δ *ulp2*Δ double mutant is suppressed by *ulp1-C376* (L) that is no longer tethered to the nuclear pore complex, and by the deletion of the STUbL subunit Slx5 (M).
